# Modeling neuron–glia interactions with the *Brian 2* simulator

**DOI:** 10.1101/198366

**Authors:** Marcel Stimberg, Dan F. M. Goodman, Romain Brette, Maurizio De Pittà

## Abstract

Despite compelling evidence that glial cells could crucially regulate neural network activity, the vast majority of available neural simulators ignores the possible contribution of glia to neuronal physiology. Here, we show how to model glial physiology and neuron-glia interactions in the *Brian 2* simulator. *Brian 2* offers facilities to explicitly describe any model in mathematical terms with limited and simple simulator-specific syntax, automatically generating high-performance code from the user-provided descriptions. The flexibility of this approach allows us to model not only networks of neurons, but also individual glial cells, electrical coupling of glial cells, and the interaction between glial cells and synapses. We therefore conclude that *Brian 2* provides an ideal platform to efficiently simulate glial physiology, and specifically, the influence of astrocytes on neural activity.

## 1. Introduction

### 1.1 Challenges in modeling of neuron–glia interactions

Computational modeling is an important part of modern neuroscience research (Abbott, 2008). Neuronal models are available at many scales of investigation (Dayan and Abbott, 2001): highly detailed multi-compartmental models describe the ion channels that establish the membrane potential and are responsible for action potential generation; simplified models, such as the integrate-and-fire neuron, regard action potentials as stereotypical events that are not described any further; and even more abstract models, such as Poisson neurons, only aim to capture the statistics of timing of action potentials rather than the shape or the biophysical mechanism of their generation. Such neuronal models can be readily simulated by a wide range of simulator packages for neuronal simulations (Brette et al., 2007), given that they are well-established and fairly standardized and thus are often provided by libraries built in these packages (but see Brette, 2012).

The same does not hold true for glial cell models (De Pittà et al., 2013). Despite compelling emerging evidence that glial cells could crucially regulate neural network activity (Poskanzer and Yuste, 2016) and plasticity (De Pittà et al., 2016), the vast majority of available neuronal models completely ignores the possible contribution of glia to neuronal physiology. Arguably, one of the reasons for this is that standard glia models are yet to be defined (Chapter 1), and thus simulator packages generally do not provide models of glial cells as part of their libraries.

Although several popular neural simulators, such as NEURON (Carnevale and Hines, 2006) or NEST (Gewaltig and Diesmann, 2007), allow for extending their built-in library with user-defined glia models, this is generally not straightforward as it requires specific programming skills in a low-level language. Additionally, it usually involves an additional step of compilation and integration into the simulator every time the library is changed. This may ultimately discourage research that involves adding or modifying a glia model in a simulator, since iterative improvements are very inconvenient. As a consequence, computational studies that model glia either use a standard simulator such as NEURON but are limited in usability by the specific choice made for the glial model which cannot easily be modified by the user (Aleksin et al., 2017), or use custom code written in a general-purpose language such as MATLAB (De Pittà et al., 2011; Naeem et al., 2015; Wade et al., 2011) or lower-level languages such as C/C++ (Nadkarni et al., 2008; Volman et al., 2007), in turn suffering from reduced accessibility to researchers with less technical experience. Overall, both scenarios raise potential issues in terms of portability, reproducibility and correctness of the code which are detrimental to model sharing and dissemination (Cannon et al., 2007).

In contrast to most other simulators, the *Brian* simulator (Goodman and Brette, 2008; Goodman et al., 2009) was created with the aim to ease definition and portability of new models. In its latest version, *Brian 2*, this flexibility is extended to and combined with a generic approach for code generation that allows high-performance simulations (Goodman, 2010; Stimberg et al., 2014). This chapter explores these aspects, elucidating several advantages that should encourage researchers to use the *Brian 2* simulator to model glia in their work.

### 1.2 The Brian simulator

The *Brian* simulator, created in 2008, is provided as a package for the Python programming language. All aspects of the model can be defined in a single Python script and are made explicit: rather than relying on predefined “black-box” models, models can be readily and flexibly described in mathematical terms by differential equations for continuous dynamics and a series of update statements for discrete events (Brette, 2012; Stimberg et al., 2014). This allows high code readability and flexibility as the user can freely change details of the model’s equations which are written in mathematical notation with only very little *Brian*-specific syntax. Furthermore, because the description of the model is explicit, all model details are unambiguously defined and appear in the main simulation description file.

In line with the principle of readability and simplicity, *Brian 2* also comes with a system for the use of physical units. For example, it allows the user to directly specify a parameter in μm/s units, by multiplying its value by umolar/second. *Brian 2* also checks the consistency of all specified units across all assignments, statements, and equations and issues an error if there is a mismatch.

Brian is open source and freely distributed under the GPL-compatible CeCILL v2.1 license. For more information see http://briansimulator.org and http://brian2.readthedocs.io.

### 1.3 Modeling strategy

In the following, we focus on astrocytes and their interactions with synapses, but the modeling arguments and code design principles that we present are of general validity and could be used to also model other glial cells, such as microglia, oligodendrocytes or reactive astrocytes. Computational modeling of neuron–astrocyte signaling has previously been tackled both on the microscopic (molecular) level and the macroscopic (network) scale. On the microscopic level, the MCell simulator (Stiles et al., 2001) has been used to investigate specific astrocytic signals impinging on synaptic elements (Beenhakker and Huguenard, 2010). On the network level, the NEURON-based ARACHNE platform to study astrocyte functions in neural network physiology is available, but only considers astrocyte-mediated ‘volume-transmitted’ extracellular signaling (Aleksin et al., 2017; Savtchenko and Rusakov, 2014).

In the following modeling section we pursue instead a mixed strategy, considering models of astrocytes and of their interactions with synapses that lump both microscopic and macroscopic aspects. Based on this approach, we show how *Brian 2* can be used to create a network of neurons and synapses connected with a network of astrocytes that sense synaptic activity and modulate it in turn, starting from a molecular-level description of astrocytic signaling. We do so by first introducing a simple network model of only neurons and synapses (Sections 2.2 and 2.3). Then, we present modeling of individual astrocytes that respond to synaptic activation by intracellular calcium signaling (Section 2.4) and release gliotransmitters that could modulate synaptic transmission (Sections 2.5 and 2.6). Next, we discuss signaling between astrocytes in a network (Section 2.7) and finally we combine all these aspects in a recurrent network of interacting neurons, synapses, and astrocytes (Section 2.8).

For the sake of brevity, we only show excerpts of the *Brian 2* code describing the models used in the presented simulations. The full code (see also Appendix B), including the code for recording and analyzing the results, as well as plotting the figures of this chapter can be downloaded from https://github.com/mdepitta/comp-glia-book. It is also part of the *Brian 2* documentation at http://brian2.readthedocs.io and available in ModelDB (McDougal et al., 2017) at http://modeldb.yale.edu/233393.

## 2 Modeling of neuron–glia network interactions with *Brian 2*

### 2.1 General approach

In *Brian 2*, models of neurons, synapses, and astrocytes are defined by a set of state variables, e.g. the neuron’s membrane potential, the synaptic conductances, or the astrocyte’s intracellular calcium, and a description of their evolution over time. This description takes the form of ordinary differential equations (ODEs) for the continuous temporal dynamics between “events”, e.g. the membrane potential, postsynaptic conductances or astrocytic intracellular calcium concentration between action potentials. Discontinuities in the dynamics of state variables triggered by events such as the crossing of the firing threshold by a neuron’s membrane potential, or the arrival of an action potential at a presynaptic terminal, are described by a set of statements that update these variables.

Groups of elements that share the same description of their dynamics are represented by a single object. Thus, for example, groups of neurons, synapses, and astrocytes would each be represented by an object. However, these groups can be heterogeneous. For example, neurons or astrocytes in the same group can be stimulated by different synapses, or synapses can be characterized by different cellular parameters. State variables are updated according to the dynamics specified by the user. If the evolution of a state variable is of interest for the purpose of analysis or visualization, the user can record it by a “monitor” object either for all elements of a group or for a subset thereof.

Once all elements of a model have been specified, including initial values, constants, and model parameters, the simulation can be launched. This is done by calling the run function with a parameter that specifies the desired total simulated time in biological time units (e.g. second). Upon completion of the run call, the user can analyze the simulation results either by accessing the final values of state variables (for example to analyze the synaptic weight distribution at the end of a simulated plasticity-inducing protocol), or by accessing values stored in a monitor. All these values are readily accessible as NumPy arrays (van der Walt et al., 2011) and can be stored, analyzed and displayed by standard tools. Since simulation results are annotated with physical units, plotting them in a specific scale can easily be done by dividing them by those units. For example, when the state variable v (“membrane potential”) of a group of neurons has been recorded by monitor=StateMonitor(neurons, ‘v’), the first neuron’s membrane potential may be shown in mV as function of time in ms using Matplotlib’s (Hunter, 2007) plot function by plot(monitor.t/ms, monitor[0].v/mV).

### 2.2 Neurons

In *Brian 2*, neurons are represented by objects of the NeuronGroup class (Figure 1A). Each NeuronGroup object models the activity of a group of neurons with identical dynamics, i.e. neurons whose state variables evolve according to the same differential equations. Consider, for example, the simple model of an integrate-and-fire neuron with conductance-based excitatory (*g*_*e*_) and inhibitory synapses (*g*_*i*_) and a constant input current *I*_*ex*_ (Dayan and Abbott, 2001), whose equations are

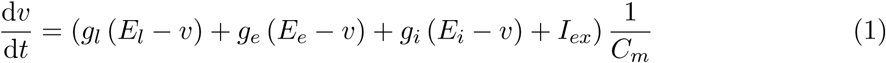

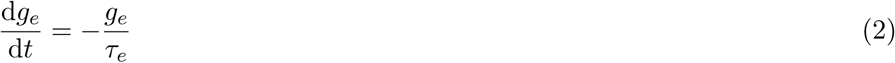

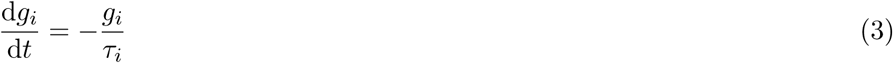

**Figure 1.**
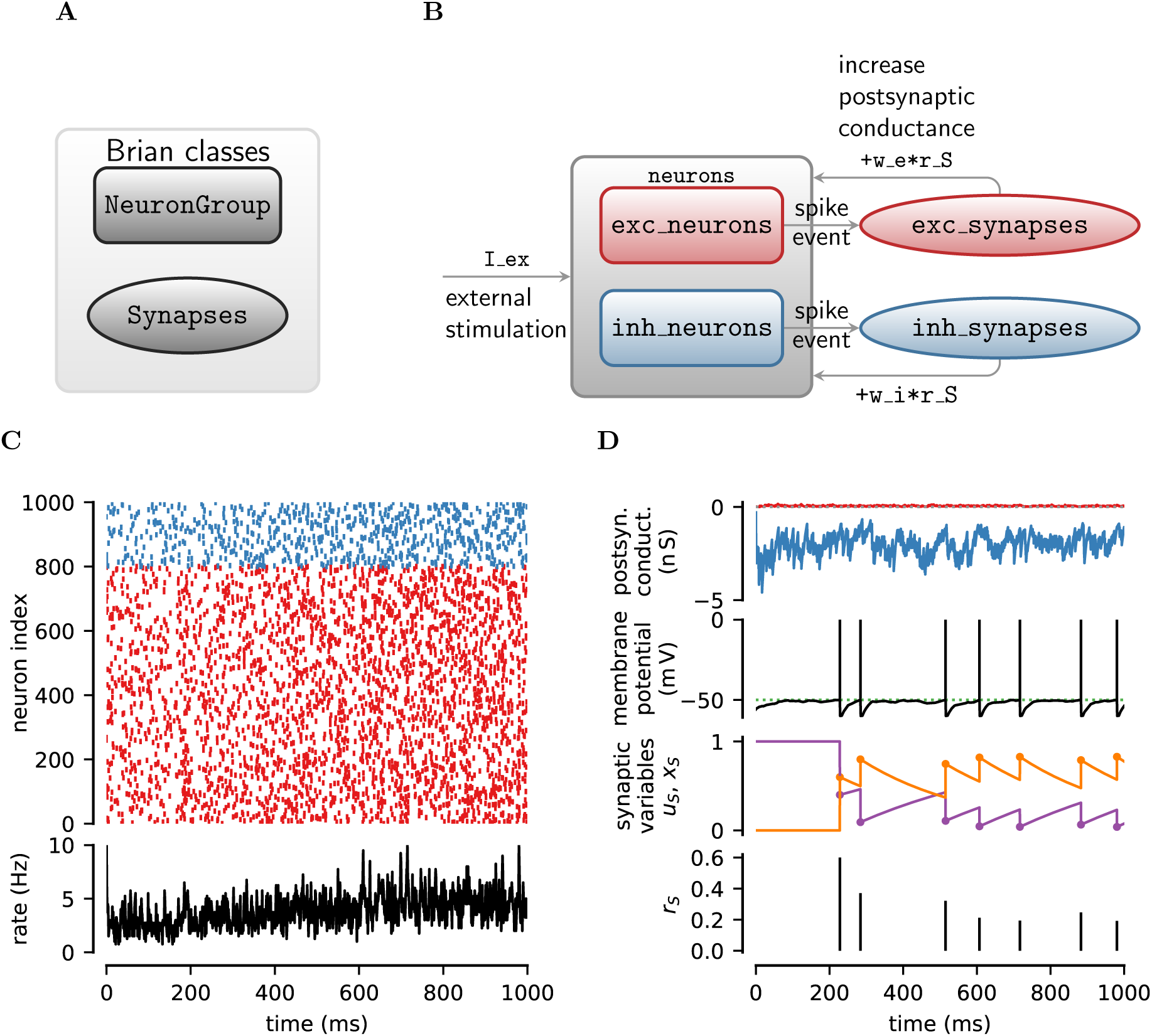
Modeling of neurons and synapses. **A** In *Brian 2* neurons and synapses are modeled by two distinct classes, `NeuronGroup and Synapses`. As a convention, in all the schemes presented in this chapter, `NeuronGroup` objects are shown as *rectangles*, whereas Synapses objects are displayed as *ellipses*. **B** In a classic balanced network model (Brunel, 2000), neurons are separated into excitatory `(exc_neurons)` and inhibitory ones `(inh_neurons)`, being recurrently connected both by excitatory `(exc_synapses)` and by inhibitory synapses (inh_synapses). **C** Raster plot of the firing activity of 25% out of all excitatory (*red*) and inhibitory neurons (*blue*) of the network in panel **B** and associated network-averaged firing rate (computed in 1 ms time bins). **D** Asynchronous network activity coexists with large fluctuations in postsynaptic excitatory (*red traces*) and inhibitory conductances (*blue traces*) and relatively sporadic firing by individual neurons (*green dotted line*: firing threshold, V_th). Timing of incoming presynaptic action potentials also shapes the dynamics of synaptic transmission by short-term synaptic plasticity. Synaptic release of neurotransmitter (*r*_*S*_) is not fixed, but rather varies at each action potential, depending on the history of synaptic activity reflected in the values of the synaptic state variables *u*_*S*_ (*orange*) and *x*_*S*_ (*purple*) at the action potential instant. Postsynaptic conductances and membrane potential are shown for neuron 50 from the raster plot. Displayed synaptic variables are from one excitatory synapse made by this neuron. Model parameters as in Table C.1 and in addition, *I*_*ex*_ = 150 pA.

**C.1.** Neurons and synapses

C.1 Neurons and synapses
| Symbol | Name in code | Value | Units | Description |
| --- | --- | --- | --- | --- |
| *Neuron parameters* |  |  |  |  |
| $C_m$ | C_m | 198 | pF | Membrane capacitance |
| $E_l$ | E_l | -60 | mV | Leak reversal potential |
| $g_l$ | g_l | 9.99 | nS | Leak conductance |
| $I_{ex}$ | I_ex | † | pA | External current |
| $V_r$ | V_r | -60 | mV | Reset potential |
| $V_\theta$ | V_th | -50 | mV | Firing threshold |
| $\tau_r$ | tau_r | 5 | ms | Refractory period |
| *Synapse parameters* |  |  |  |  |
| $\Omega_d$ | Omega_d | 2 | s<sup>-1</sup> | Synaptic depression rate |
| $\Omega_f$ | Omega_f | 3.33 | s<sup>-1</sup> | Synaptic facilitation rate |
| $U_0^*$ ($U_0$) | U_0__star (U_0) | 0.6 | – | Resting synaptic release probability |
| $Y_T$ | Y_T | 500 | mM | Total vesicular neurotransmitter concentration |
| $\rho_c$ | rho_c | 0.005 | – | Synaptic vesicle-to-extracellular space volume ratio |
| $\Omega_c$ | Omega_c | 40 | s<sup>-1</sup> | Neurotransmitter clearance rate |
| $w_e$ | w_e | 50 | pS | Excitatory synaptic conductance |
| $w_i$ | w_i | 1 | nS | Inhibitory synaptic conductance |
| $\tau_e$ | tau_e | 5 | ms | Excitatory synaptic time constant |
| $\tau_i$ | tau_i | 10 | ms | Inhibitory synaptic time constant |
| $E_e$ | E_e | 0 | mV | Excitatory synaptic reversal potential |
| $E_i$ | E_i | -80 | mV | Inhibitory synaptic reversal potential |
| *Presynaptic receptors* |  |  |  |  |
| $O_G$ | O_G | 1.5 | $\mu\text{M}^{-1} \text{s}^{-1}$ | Agonist binding (activating) rate |
| $\Omega_G$ | Omega_G | 0.5 | min<sup>-1</sup> | Agonist release (deactivating) rate |

In *Brian 2* we closely follow the above mathematical notation, defining the neuron model by a multi-line string:

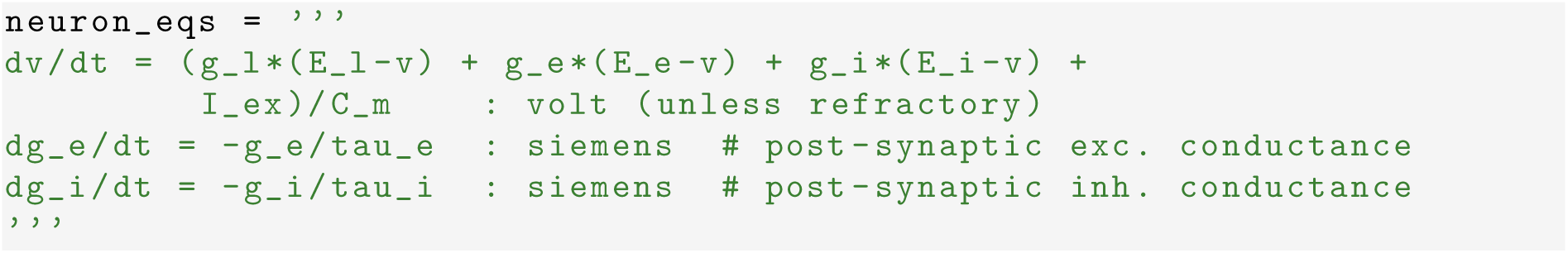

Each line of the `neuron_eqs` string defines a state variable of the model (`v, g_e or g_i`) and consists of two parts separated by a colon and an optional comment after the # sign. The part before the colon specifies the ODE for the dynamics of the state variable. The part that follows the colon specifies the physical units of this state variable and (optionally) additional information about it. In the case of the neuron’s membrane potential v for example, the additional specification (`unless refractory`) states that the differential equation for v is not to be integrated during the refractory period following the firing of an action potential by the neuron, but instead should stay at the post-spike reset value. It should be noted that the stated physical unit after the colon has to be a SI base unit, i.e. a unit such as volt, second, or siemens, and not a scaled unit such as mV, ms, or nS^1^. This is to emphasize that internally all variable values are stored as floating point numbers in the base unit. While users do not have to care about this most of the time – they set values using the unit system in whatever scale they prefer and receive values with the scaling information back – this become relevant when the unit information is stripped away, e.g. when quantities are passed through library functions that are not unit-aware.

The implementation of the neuron model in equations 1–3 refers to model parameters, namely the leak, excitatory, and inhibitory reversal potentials *E*_*l*_, *E*_*e*_ and *E*_*i*_; the constant input current *I*_*ex*_; the membrane capacitance *C*_*m*_; and the time constants of excitatory and inhibitory synaptic inputs, *τ_e_* and *τ_i_*. Here, these parameters are taken to be equal for all simulated neurons and can be defined by standard Python variables (one per parameter) in the script that runs the simulation^2^. Alternatively, neuron-specific parameters can be defined by appending lines in the form of `<name>: <unit> (constant)` to the model equations.

In the scenario of a network of *N*_*e*_ excitatory and *N*_*i*_ inhibitory neurons (Figure 1B), we can then create a `NeuronGroup` object of *N*_*e*_ +*N*_*i*_ neurons based on the above description, and further define the condition for firing of an action potential `(threshold)`, as well as the statement(s) (if any) to be executed after an action potential `(reset)`, and finally the refractory period (refractory), which in this case we assume to be a constant value *τ_r_* (defined along with the other model parameters):

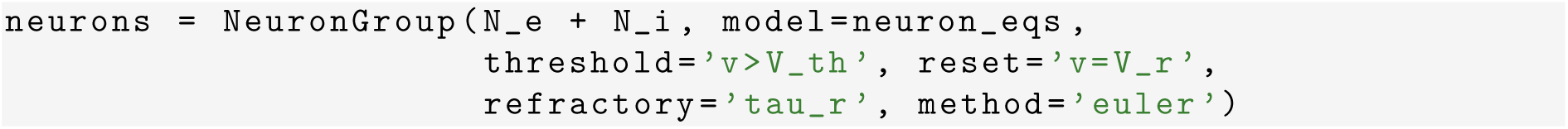

The model’s state variables are exposed as attributes of the neurons object so that, for example, the membrane potential can be accessed by neurons.v, and their initial value can be assigned in the same way. Although all neurons in our example are described by the same equations, the initial values of their variables can be different. Here, we set these latter to random values using string expressions that are executed via the code generation facilities provided by *Brian 2* (Stimberg et al., 2014) and refer to uniformly distributed random numbers between 0 and 1 using the predefined function rand(). Finally, we use Python’s slicing syntax to separate the group into subgroups of excitatory and inhibitory neurons:

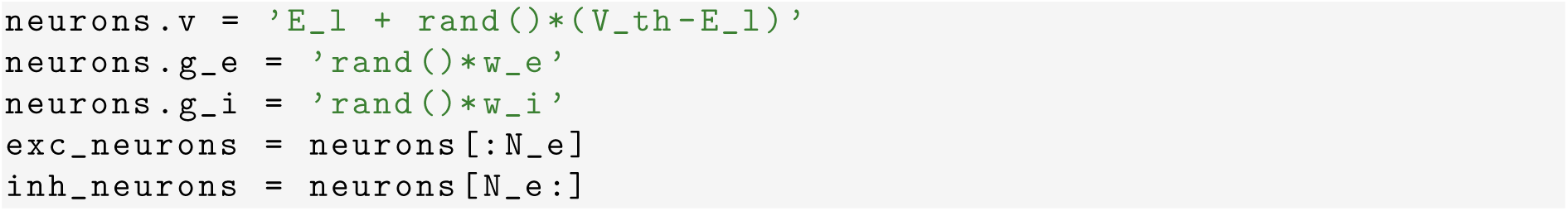

### 2.3 Synapses

In most models of neuronal systems, neurons are connected by chemical synapses that are activated by action potentials fired by presynaptic neurons. In the following, we use the phenomenological description of neocortical synapses exhibiting short-term plasticity originally introduced by Tsodyks and Markram (Tsodyks, 2005; Tsodyks et al., 1998). According to this description, synaptic release is modeled by the product of two variables *u*_*S*_ and *x*_*S*_, where *u*_*S*_ loosely relates to the neurotransmitter resources “docked” for release by the Ca^2+^ sensor for synaptic exocytosis of neurotransmitter, and *x*_*S*_ represents the fraction of total neurotransmitter available for release (Fuhrmann et al., 2002; Tsodyks, 2005). Between action potentials, *u*_*S*_ decays to 0 at rate Ω*_f_* while *x*_*S*_ recovers to 1 at rate Ω*_d_*, i.e.

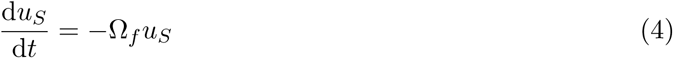

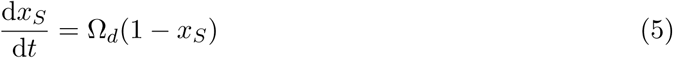

The arrival of an action potential triggers calcium influx at the presynaptic terminal, which moves a fraction *U*_0_ of the neurotransmitter resources not scheduled for release (1 *- u_S_*) to the readily-releasable “docked” state (*u*_*S*_). Subsequently, a fraction *u*_*S*_ of the available neurotransmitter resources is released as *r*_*S*_ while *x*_*S*_ is reduced by the same amount, that is

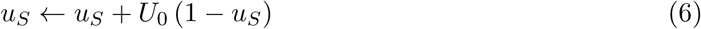

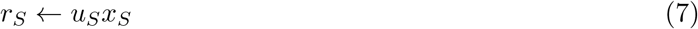

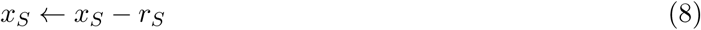

In *Brian 2*, connections between neurons are modeled by objects of the `Synapses` class (Figure 1A). Analogously to neurons of a NeuronGroup, we define each synapse’s state variables *x*_*S*_ and *u*_*S*_ by a multi-line string:

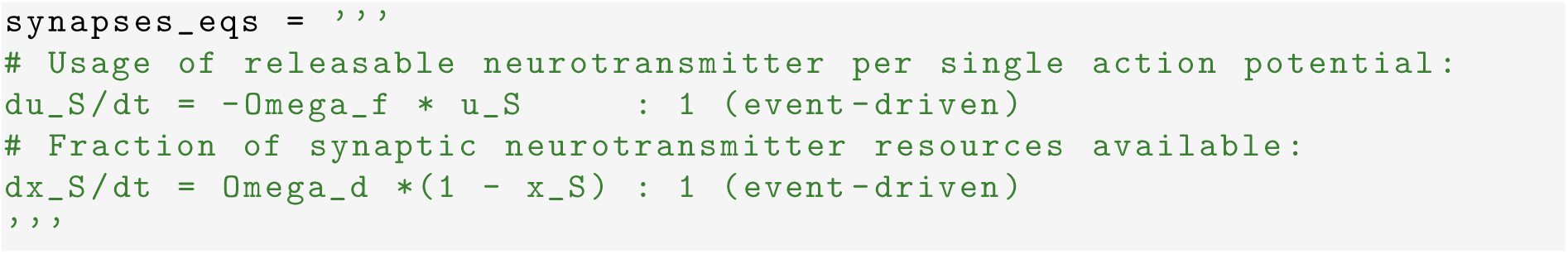

Synaptic equations are specified to be `(event-driven)` because we only need values of synaptic variables when an action potential arrives presynaptically. This tells *Brian 2* to update synaptic variables only at new incoming action potentials using the analytical solution of their differential equations and the time interval since the last update. This is possible because the synapse’s equations 4 and 5 are linear so that their analytical solution is known, allowing to simulate a large number of synapses efficiently. We also note that in our model implementation, synaptic equations do not include postsynaptic conductances as they were previously defined in the neuronal equations. This allows storing and updating synaptic conductances only once per neuron rather than once per synapse and is mathematically equivalent due to the assumed linear summation of postsynaptic conductances.

Discrete changes of synaptic variables on arrival of an action potential can be implemented by a series of statements in a multi-line string such as

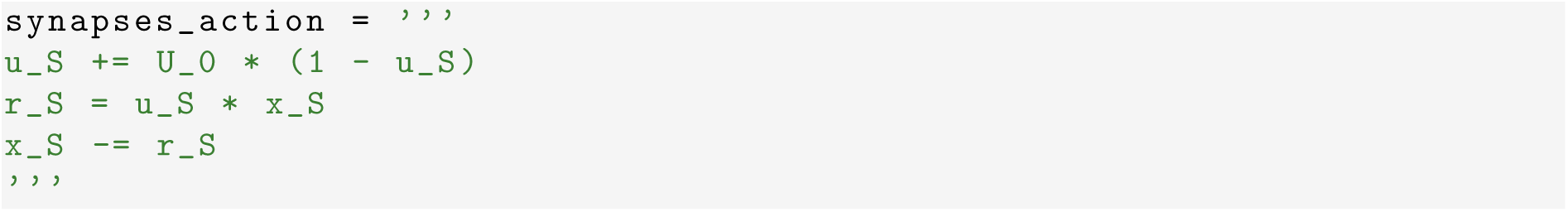

In addition, excitatory (inhibitory) synapses will increase the excitatory (inhibitory) conductance in the postsynaptic cell whenever a presynaptic action potential arrives (Figure 1B), i.e.

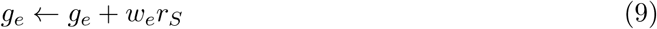

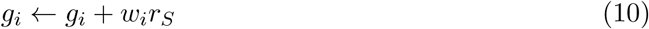

This is done by extending synapses_action by an update assignment ‘+=’ of the respective postsynaptic conductance, identified by the post suffix, i.e. g_e_post and g_i_post. Combining all of this together, we create two types of synapses, respectively originating from excitatory and inhibitory neurons, i.e.

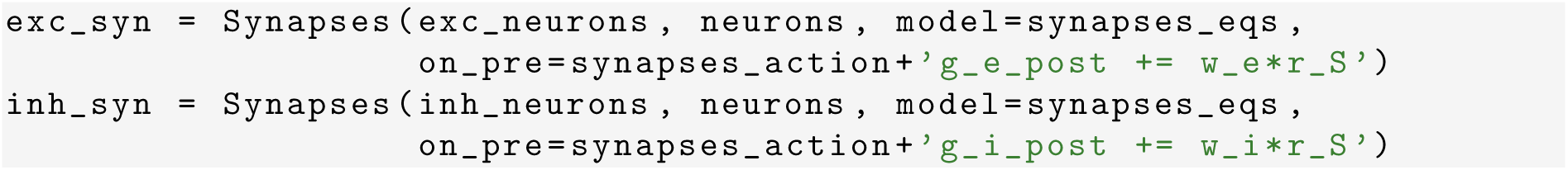

where the on_pre keyword argument denotes that the series of statements should be executed on arrival of a presynaptic action potential.

It must be stressed that the above code only defines the synaptic model in *Brian 2*, but not the connectivity. To create synapses, we have to specify what source–target neuron pairs should be connected together out of all the possible pairs specified by the first two input arguments in the `Synapses` initializer. One way to do this is by specifying a logical condition on neuronal connectivity and, optionally, a connection probability, provided as arguments to the connect method of the `Synapses` object (Stimberg et al., 2014). Here, we want to connect all possible neuron pairs with a probability of 20% for each pair for inhibitory neurons, and 5% for excitatory neurons. Thus we do not set any condition to be fulfilled and only specify a probability for all possible connection pairs:

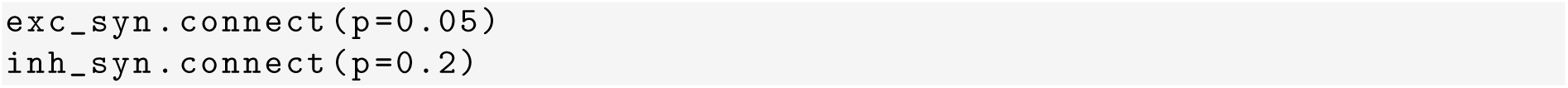

For parameters values such as those in Table C.1, simulation of the resulting network reveals the emergence of characteristic asynchronous neuronal firing activity (Brunel, 2000), as evidenced in Figure 1C by the *top left* raster plot of the firing activity of 25% of the neurons in the network. The network-averaged firing rate associated with this raster plot indeed presents noisy dynamics (Figure 1C, *bottom panel*) that coexists with large fluctuations of postsynaptic excitatory (*red traces*) and inhibitory conductances (*blue traces*) and sporadic firing of individual neurons (Figure 1D, *top panels*). Consideration of a sample excitatory synapse of the network allows appreciating how synaptic dynamics is modulated by short-term plasticity (Figure 1D, *bottom panels*). The state variables *u*_*S*_ and *x*_*S*_ associated with the sample synapse evolve in a characteristic exponential fashion intermingled with discontinuities triggered by action potentials arriving at the presynaptic terminal. The resulting values define the amount of synaptic neurotransmitter released by the synapse (*r*_*S*_), setting how effectively each action potential is transmitted to the postsynaptic neuron.

### 2.4 Astrocytes

Intracellular Ca^2+^ concentration is unanimously regarded as a prominent readout signal of astrocyte activity (Zorec et al., 2012). Although astrocytic intracellular Ca^2+^ can be regulated by multiple mechanisms, Ca^2+^-induced Ca^2+^ release (CICR) from the astrocyte’s endoplasmic reticulum (ER) appears to be one of the main mechanisms to regularly occur in the healthy brain (Nimmerjahn, 2009). Recall from Chapter 5 that astrocytic CICR is triggered by the intracellular second messenger inositol 1,4,5-trisphosphate (IP_3_), which is produced upon astrocyte activation, and can be described, in one of its simplest formulation (De Pittà et al., 2008; Falcke, 2004), by two ordinary differential equations in the Hodgkin-Huxley form (Li and Rinzel, 1994). The first equation is a mass balance for Ca^2+^ (*C*) in terms of three fluxes *J*_*r*_, *J*_*l*_, *J*_*p*_ which respectively denote CICR (*J*_*r*_), Ca^2+^ leak from the ER (*J*_*l*_), and Ca^2+^ uptake from the cytosol back to the ER by Ca^2+^/ATPase pumps (*J*_*p*_). The second equation is for the gating variable (*h*) of de-inactivation of the channels that are responsible for CICR. These channels are inside the astrocyte, on the membrane that separates the ER Ca^2+^-rich stores from the cell’s cytosol, and are nonlinearly gated by both IP_3_ (*I*) and Ca^2+^. This leads to the well-known two-equation model originally introduced by Li and Rinzel (1994):

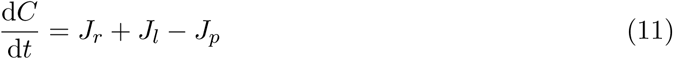

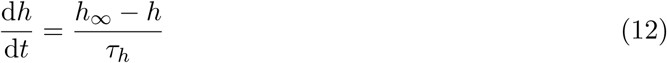

where

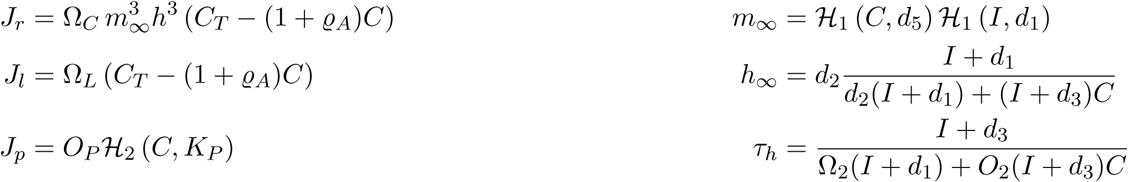

and *H* denotes the sigmoidal (Hill) function with *ℋ_n_* (*x, K*) = *x*^*n*^/ (*x*^*n*^ + *K*^*n*^).

For the sake of completeness, we also want to consider the stochastic opening and closing process of CICR-mediating channels which, as discussed in Chapter 4, can be mimicked by introducing a white noise term *ξ*(*t*) into the equation governing the dynamics of the gating variable *h*, so that equation 12 becomes (Shuai and Jung, 2002):

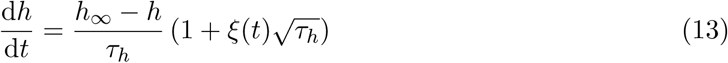

Astrocytic IP_3_ is regulated by the complex Ca^2+^-modulated interplay of enzymatic production by phospholipase C*β* (*J*_*β*_) and C*δ* (*J*_*δ*_) and degradation by IP_3_ 3-kinase (*J*_3*K*_) and inositol polyphosphatase 5-phosphatase (*J*_5*P*_) (De Pittà et al., 2009a). To reproduce experimental observations we consider two possible ways to trigger IP_3_ production. One is by synaptic stimulation of astrocytic metabotropic receptors which starts phospholipase Cβ-mediated IP_3_ production, modeled by making *J*_*β*_ proportional to the activated fraction of these receptors (denoted hereafter by Γ*_A_*). The other way is to include a further *J*_*ex*_ term for constant IP_3_ production by an exogenous source of stimulation such as, for example, IP_3_ uncaging or intracellular diflusion from subcellular regions far from the CICR site (Goldberg et al., 2010). In conclusion, IP_3_ dynamics evolves according to the mass balance equation (De Pittà et al., 2009a)

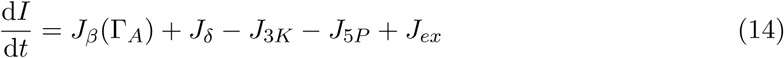

With

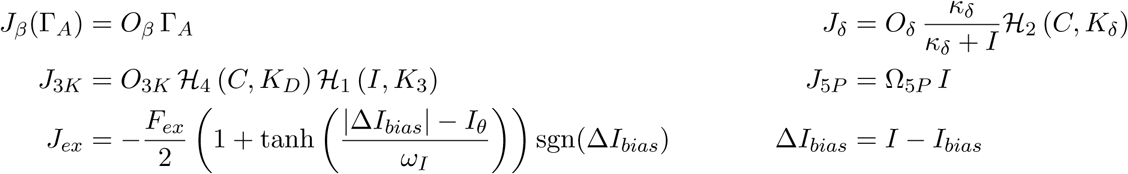

where the fraction of activated astrocyte receptors γ*_A_* depends on the neurotransmitter concentration in the periastrocytic space *Y*_*S*_, and is given by the further equation (Wallach et al., 2014) (Chapter 5)

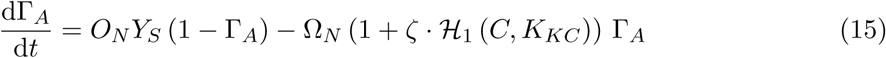

For a concise description of the meaning of all the model’s parameters in the above equations, see Table C.2 in the Appendix.

**C.2.** Astrocytes

C.2 Astrocytes
| Symbol | Name in code | Value | Units | Description |
| --- | --- | --- | --- | --- |
| *Calcium-induced $\text{Ca}^{2+}$ release* |  |  |  |  |
| $C_T$ | C_T | 2 | $\mu\text{M}$ | Total cell free $\text{Ca}^{2+}$ content |
| $\rho_A$ | rho_A | 0.18 | – | ER-to-cytoplasm volume ratio |
| $d_1$ | d_1 | 0.13 | $\mu\text{M}$ | $\text{IP}_3$ dissociation constant |
| $d_2$ | d_2 | 1.05 | $\mu\text{M}$ | $\text{Ca}^{2+}$ inactivation dissociation constant |
| $d_3$ | d_3 | 0.9434 | $\mu\text{M}$ | $\text{IP}_3$ dissociation constant |
| $d_5$ | d_5 | 0.08 | $\mu\text{M}$ | $\text{Ca}^{2+}$ activation dissociation constant |
| $O_2$ | O_2 | 0.2 | $\mu\text{M s}^{-1}$ | $\text{IP}_3\text{R}$ binding rate for $\text{Ca}^{2+}$ inhibition |
| $\Omega_C$ | Omega_C | 6 | $\text{s}^{-1}$ | Maximal rate of $\text{Ca}^{2+}$ release by $\text{IP}_3\text{Rs}$ |
| $\Omega_L$ | Omega_L | 0.1 | $\text{s}^{-1}$ | Maximal rate of $\text{Ca}^{2+}$ leak from the ER |
| $O_P$ | O_P | 0.9 | $\mu\text{M s}^{-1}$ | Maximal $\text{Ca}^{2+}$ uptake rate by SERCAs |
| $K_P$ | K_P | 0.05 | $\mu\text{M}$ | $\text{Ca}^{2+}$ affinity of SERCAs |
| *$\text{IP}_3$ signaling* |  |  |  |  |
| $O_\beta$ | O_beta | † | $\mu\text{M s}^{-1}$ | Maximal rate of $\text{IP}_3$ production by $\text{PLC}\beta$ |
| $O_\delta$ | O_delta | 0.6 | $\mu\text{M s}^{-1}$ | Maximal rate of $\text{IP}_3$ production by $\text{PLC}\delta$ |
| $\kappa_\delta$ | kappa_delta | 1.5 | $\mu\text{M}$ | Inhibition constant of $\text{PLC}\delta$ by $\text{IP}_3$ |
| $K_\delta$ | K_delta | 0.1 | $\mu\text{M}$ | $\text{Ca}^{2+}$ affinity of $\text{PLC}\delta$ |
| $O_{3K}$ | O_3K | 4.5 | $\mu\text{M s}^{-1}$ | Maximal rate of $\text{IP}_3$ degradation by $\text{IP}_3\text{-3K}$ |
| $K_{3K}$ | K_3K | 1 | $\mu\text{M}$ | $\text{IP}_3$ affinity of $\text{IP}_3\text{-3K}$ |
| $K_D$ | K_D | 0.7 | $\mu\text{M}$ | $\text{Ca}^{2+}$ affinity of $\text{IP}_3\text{-3K}$ |
| $\Omega_{5P}$ | Omega_5P | 0.05 | $\text{s}^{-1}$ | Maximal rate of $\text{IP}_3$ degradation by $\text{IP 5P}$ |
| *Metabotropic receptor kinetics* |  |  |  |  |
| $O_N$ | O_N | 0.3 | $\mu\text{M}^{-1} \text{s}^{-1}$ | Agonist binding rate |
| $\Omega_N$ | Omega_N | 0.5 | $\text{s}^{-1}$ | Maximal inactivation rate |
| $K_{KC}$ | K_KC | 0.5 | $\mu\text{M}$ | $\text{Ca}^{2+}$ affinity of PKC |
| $\zeta$ | zeta | 10 | – | Maximal reduction of receptor affinity by PKC |
| *$\text{IP}_3$ stimulation & diffusion* |  |  |  |  |
| $F_{ex}$ | F_ex | 2 | $\mu\text{M s}^{-1}$ | Maximal exogenous $\text{IP}_3$ flow |
| $I_{bias}$ | I_bias | † | $\mu\text{M}$ | External $\text{IP}_3$ drive |
| $F$ | F | 0.09 | $\mu\text{M s}^{-1}$ | GJC $\text{IP}_3$ permeability |
| $I_\theta$ | I_Theta | 0.3 | $\mu\text{M}$ | Threshold gradient for $\text{IP}_3$ diffusion |
| $\omega_I$ | omega_I | 0.05 | $\mu\text{M}$ | Scaling factor of diffusion |
| *Gliotransmission* |  |  |  |  |
| $C_\theta$ | C_Theta | 0.5 | $\mu\text{M}$ | $\text{Ca}^{2+}$ threshold for exocytosis |
| $G_T$ | G_T | 200 | $\text{mM}$ | Total vesicular gliotransmitter concentration |
| $\Omega_A$ | Omega_A | 0.6 | $\text{s}^{-1}$ | Gliotransmitter recycling rate |
| $U_A$ | U_A | 0.6 | – | Gliotransmitter release probability |
| $\rho_e$ | rho_e | 0.00065 | – | Astrocytic vesicle-to-extracellular volume ratio |
| $\Omega_e$ | Omega_e | 60 | $\text{s}^{-1}$ | Gliotransmitter clearance rate |
| $\alpha$ | alpha | 0 | – | Gliotransmission nature |

Dynamics of the astrocyte’s state variables γ*_A_, I, C, h* are governed by ODEs akin to neuronal state variables, although on a longer time scale (De Pittà et al., 2009b). Accordingly, they can be implemented by a `NeuronGroup` object, exactly in the same way as neuronal variables. In this fashion, the following code exemplifies how we can create two astrocytes with dynamics governed by the above equations:

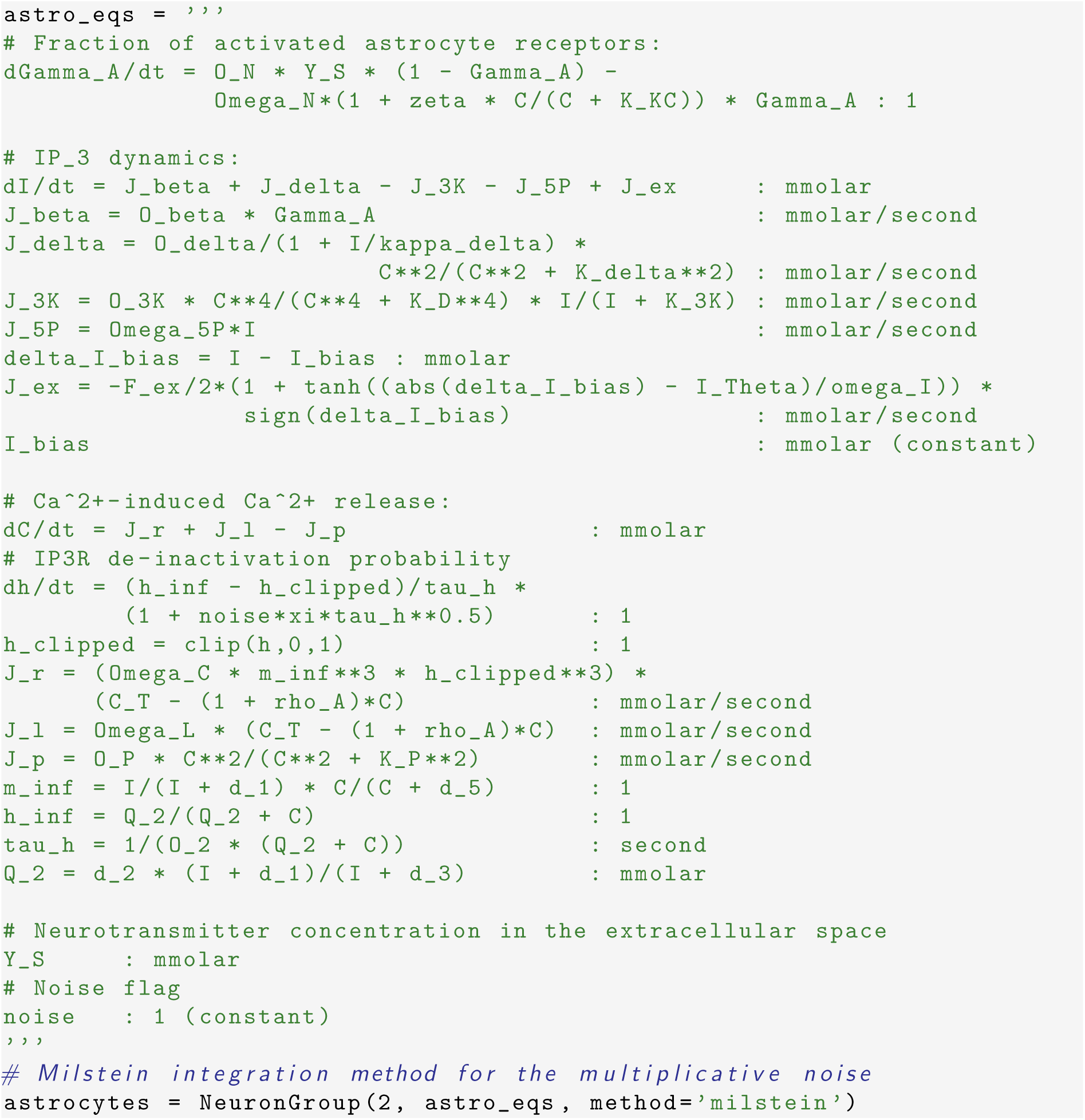

In the above *Brian 2* code, the multi-line astro_eqs string defines our astrocyte model. The white noise term *ξ* in equation 13 is implemented by the special symbol xi (in units of s^−1/2^) which is a predefined symbol in *Brian 2*. Also note that as a gating variable, *h* has to be bound between 0 and 1, which is not guaranteed by the (approximated) nature of equation 13. We therefore restrict h to this interval by replacing it by `h_clipped=clip(h,0,1)` on the right hand side of the ODE for h and in the formula of `J_r` (Shuai and Jung, 2002). The noise term in equation 13 is multiplicative (*ξ*(*t*) multiplies the gating variable *h*), we therefore have to use a numerical integration method that can handle multiplicative stochastic differential equations (under the Stratonovich interpretation, cf. Kloeden and Platen, 1992). We do this by specifying method=’milstein’ as an argument to the `NeuronGroup` initializer, leading to the use of the Milstein method for integration (Kloeden and Platen, 1992; Mil’shtejn, 1975, also see details on example 2 gchi astrocyte.py in Appendix B).

The above model also defines three astrocyte-specific variables that are not defined by equations: `I_bias, Y_S`, and noise. `I_bias` and noise are constant over time, but `Y_S`, the concentration of synaptically-released neurotransmitter in the extracellular space around astrocytic receptors, i.e. *Y*_*S*_ in equation 15, depends on synaptic activity that changes and therefore does not have the (constant) flag. We will define how it gets linked to the synaptic activity further below.

In this example, we want to compare two types of astrocytes, a deterministic and a stochastic one. To distinguish them, we have introduced the above-mentioned noise constant which scales the strength of the noise term in equation 13. We can therefore switch the noise term on or off, and we initialize it so that the first astrocyte is deterministic and the second is stochastic:

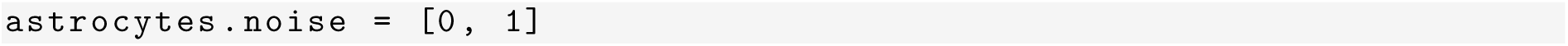

To complete our model, we also need to specify how to calculate *Y*_*S*_ in equation 15, as it is needed for the integration of the γ*_A_* state variable. For now, we are only interested in the activity of the astrocyte and how it is stimulated by synaptic neurotransmitter. Therefore, we do not take into account short-term synaptic plasticity, and rather consider a trivial synaptic model stating that *Y*_*S*_ increases by the same amount at every action potential and then decays exponentially (Dayan and Abbott, 2001):

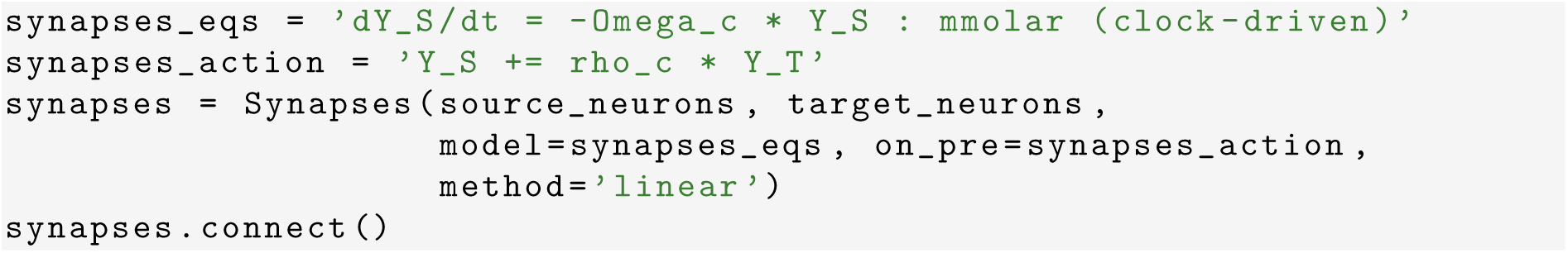

Finally we have to specify how the synapses stimulate the astrocytes. We do this by introducing a further `Synapses` object that connects our synapses (which thus represent the “presynaptic” source in our object) to the two astrocytes (hence regarded as “postsynaptic” targets) in an all-to-all fashion, which can be concisely expressed by a call to connect() without any arguments. Each astrocyte *i* senses the sum of *Y*_*S*_ across all *S*^*i*^ synapses that impinge on it. This can be mathematically expressed as 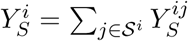, and the implementation in *Brian 2* closely follows this formulation, using the flag (summed) to denote the summing operation:

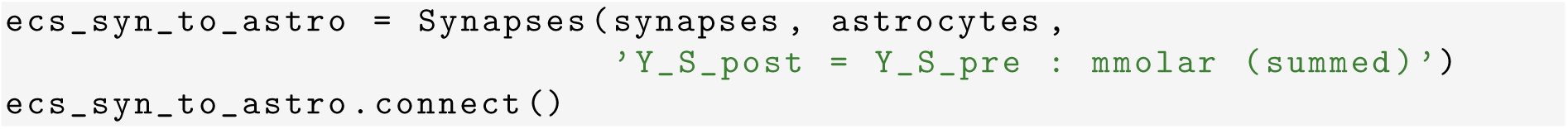

The above definition will update *Y*_*S*_ for each astrocyte at every time step so that the integration of the γ*_A_* has access to the current value at all times.

Figure 2A summarizes the design of the astrocyte model previously described, exemplifying its implementation by *Brian 2* classes originally introduced in Figure 1A. A sample simulation of this model is presented in Figure 2B, which shows the dynamics of the state variables γ*_A_, I, C* and *h* for the deterministic (*red traces*) vs. stochastic astrocyte (*blue traces*) in response to synaptic stimulation by periodic action potentials at a rate of 0.5 Hz. It may be appreciated how noisy dynamics of the gating variable *h* could dramatically alter *C* and *I* dynamics compared to the deterministic scenario. This could also impact the activated fraction of astrocytic metabotropic receptors (γ*_A_*) by the Ca^2+^-dependent Hill nonlinearity in the right hand side of equation 15, although in this example, the effect may be deemed moderate for the specific choice of values for the model’s parameters (Table C.2).

**Figure 2.**
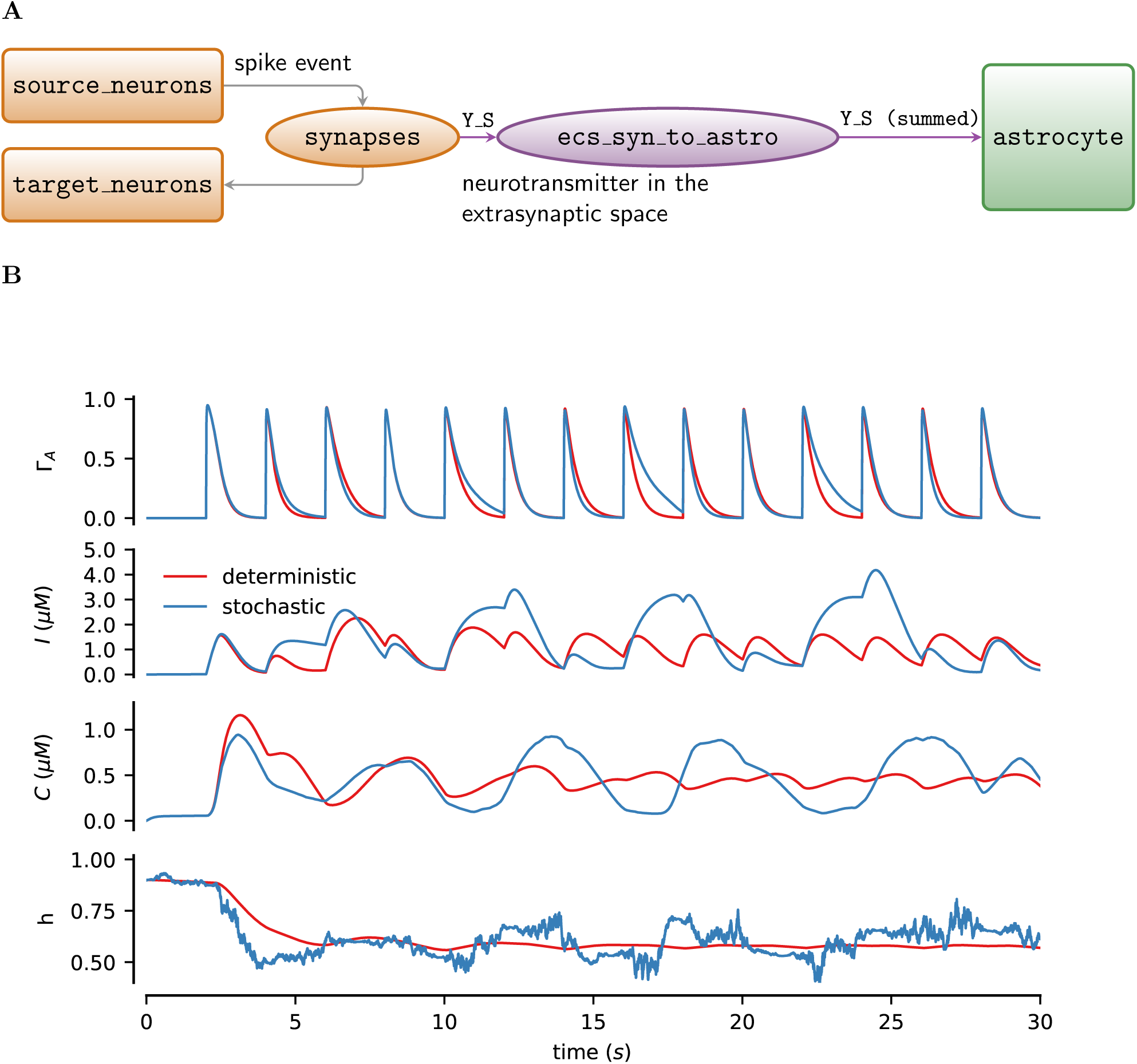
Modeling of synaptically-activated astrocytes. **A** Illustration of the design principles to implement an astrocyte stimulated by synaptic inputs in *Brian 2*. The coupling between synapses and the astrocyte is achieved by an additional `Synapses` object, labeled here as `ecs_syn_to_astro`, which feeds into the astrocyte the sums of all individual synaptic inputs. **B** Simulated dynamics of the state variables of two synaptically-activated astrocytes, one deterministic (*red traces*) and one with noise on the gating variable *h* according to equation 13 (*blue traces*). The two astrocytes shared the same synaptic input which was repeatedly triggered by action potentials arriving at a rate of 0.5 Hz. Model parameters as documented in tables in Appendix C with the exception of *ρ_c_* = 0.001; *K*_*P*_ = 0.1 μm; *O*_*β*_ = 5 μm s^−1^; *O*_*δ*_ = 0.2 μm s^−1^; Ω_5*P*_ = 0.1 s^−1^; *K*_*D*_ = 0.5 μm; *F*_*ex*_ = 0.09 μm; *K*_*δ*_ = 0.3 μm; and *I*_*bias*_ = 0.

### 2.5 Gliotransmitter release and modulation of synaptic release

Astrocytes are not only stimulated by synapses but they can also modulate them by releasing neurotransmitters (also termed “gliotransmitters” for their glial origin; Bezzi and Volterra, 2001) in a Ca^2+^-dependent fashion. This process generally requires astrocytic intracellular Ca^2+^ concentration to increase beyond a certain threshold, resulting in the release of a quantum of gliotransmitter into the periastrocytic space (De Pittà et al., 2013). In turn, released gliotransmitter molecules diffuse in the extracellular space, ultimately reaching extrasynaptic receptors found on synaptic elements belonging either to the very synapses that stimulate the astrocyte in the so-called “closed-loop” scenario of gliotransmission, or to other synapses in the case of “open-loop” gliotransmission (Araque et al., 2014). Among these targeted receptors, presynaptically-located ones, once bound by gliotransmitters, can ultimately regulate synaptic transmission through modulations of synaptic release probability (Engelman and MacDermott, 2004; Pinheiro and Mulle, 2008). In the simplest approximation, as elucidated in Chapter 12, this modulation can be modeled by treating the parameter *U*_0_ in the previously-introduced Tsodyks-Markram synapse model (equation 4) no longer as a constant, but rather as linearly dependent on the fraction γ*_S_* of activated presynaptic receptors (De Pittà et al., 2011):

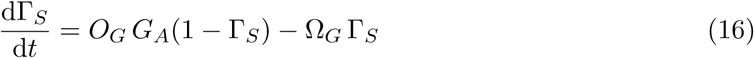

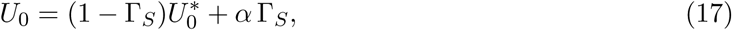

where *G_A_* denotes the gliotransmitter concentration in the extracellular space, and *α* dictates whether the effect of gliotransmitters on the synapse is to decrease 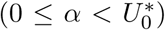 or increase neurotransmitter release 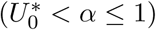 (De Pittà et al., 2011). In *Brian 2* syntax, this leads to the following synaptic equations:

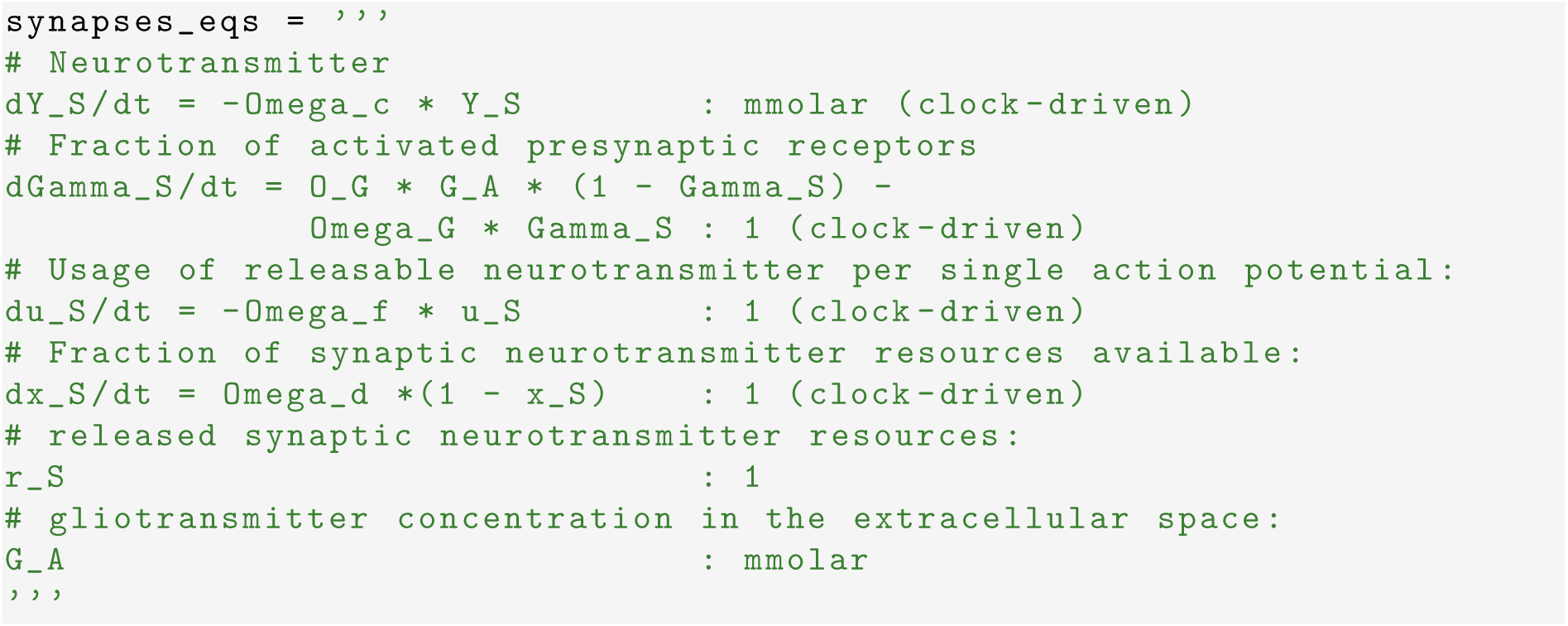

Because the value of *U*_0_ is only needed at the arrival of an action potential, there is no need to include equation 17 in the above code. Rather, we update *U*_0_ at the beginning of the statements executed by synapses upon action potential arrival, i.e.

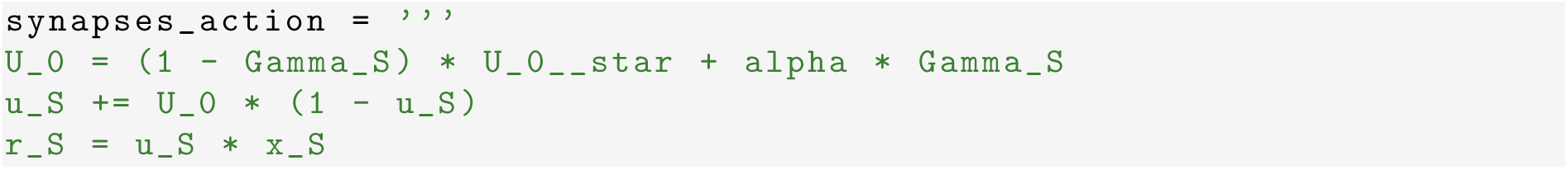

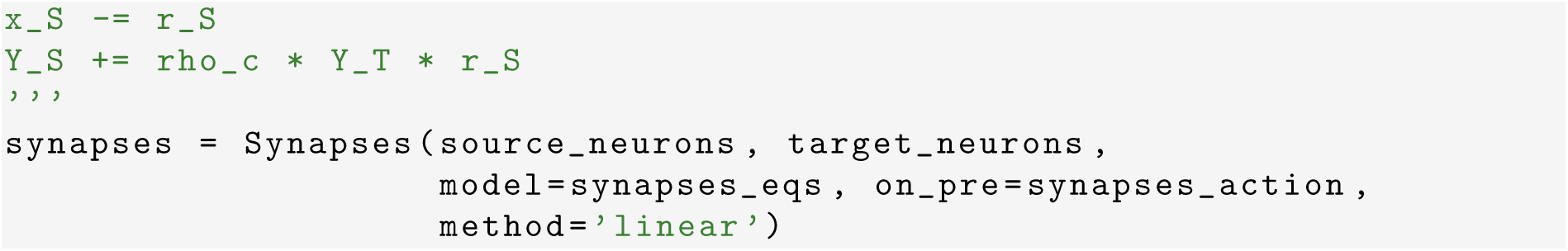

For the sake of simplicity we retained two inefficiencies in the above code which should otherwise be avoided in larger, computationally-demanding simulations. First, we used the (clock-driven) specification (instead of (event-driven)), even though synaptic state variables need only be updated on action potential arrival (Section 2.3). This allows us to directly retrieve and plot state variables at each time step without the need to manually interpolate between their values at the timing of action potentials. For the same reason, we also defined *r*_*S*_ (equation 7) as an additional state variable in the synapse model in `synapses_eqs` rather than using it as an auxiliary temporary variable in the statements of `synapses_action` as we previously did (Section 2.3). This choice allows us to easily record *r*_*S*_ by a monitor, avoiding the need to recompute it a posteriori based on the values of the other state variables.

Finally, we need to model gliotransmitter release from the astrocyte. For this, we use a similar description to that of synaptic neurotransmitter release (Chapter 13). We thus introduce a new variable *x_A_* which represents the fraction of gliotransmitter resources available for release from the astrocyte and thereby controls the value of *G_A_*. These two state variables decay as (De Pittà et al., 2011)

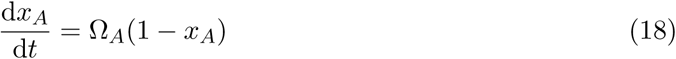

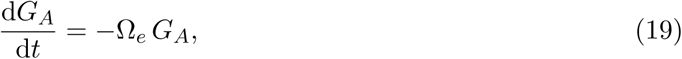

while, when gliotransmitter is released, they are updated according to

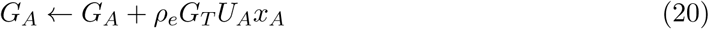

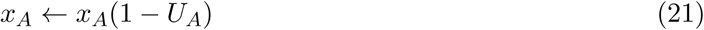

As before, we can implement the above by textual equations and statements in *Brian 2*, i.e.

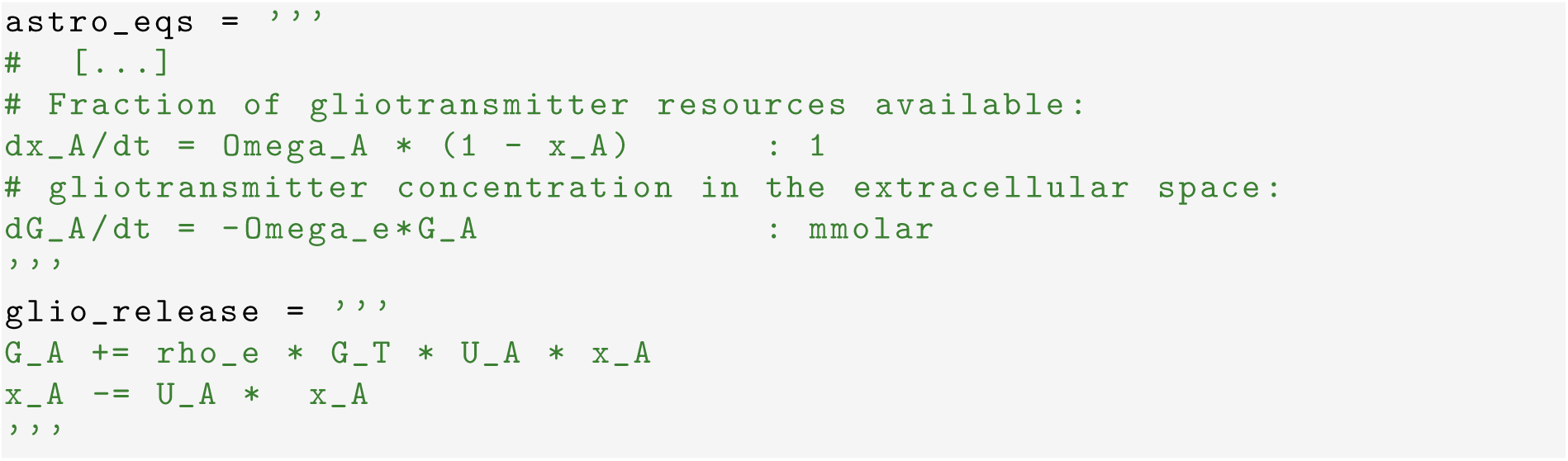

We also need to include in the astrocytic model a mechanism to time gliotransmitter release. We do this by imaging that when the astrocyte’s Ca^2+^ concentration increases beyond a threshold *C_θ_*, the astrocyte “fires” a gliotransmitter release event upon which the statements of `glio_release` are executed. In this fashion, we can define in the astrocyte’s `NeuronGroup` a threshold crossing for Ca^2+^ concentration (by the threshold keyword), upon which gliotransmitter is released, and specify by the reset keyword what to do following firing of a gliotransmitter release event by the astrocyte. Moreover, to avoid the threshold condition repeatedly triggering the release in all the time steps where the Ca^2+^ concentration is above the threshold, we use the same condition for the refractory keyword, thereby stating that, as long as the Ca^2+^ concentration is above threshold, no new event should be triggered. That is,

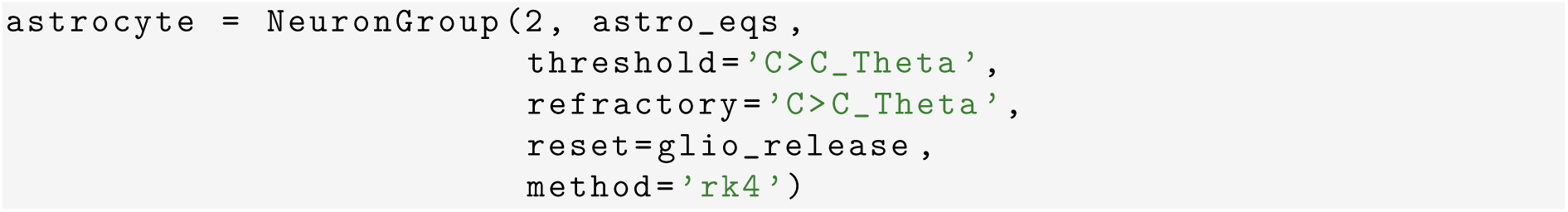

Finally, to complete our model, we have to define how synapses are modulated by the astrocytes’ gliotransmitter release. We do this defining another `Synapses` object as exemplified in Figure 3A, akin to what we did in the previous section to connect synaptic neurotransmitter release to astrocytes (Figure 2A). However, the connection between synapses and the astrocyte is now in the opposite direction, i.e astrocytic gliotransmission is upstream (namely “presynaptic”) with respect to synapses (which are thus “postsynaptic”). Hence,

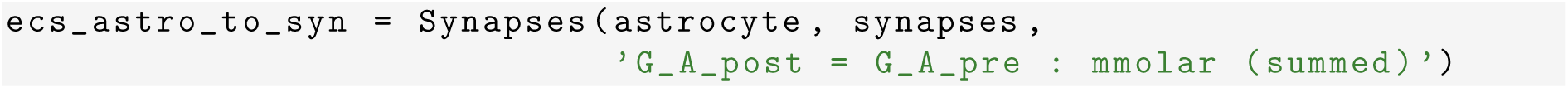

**Figure 3.**
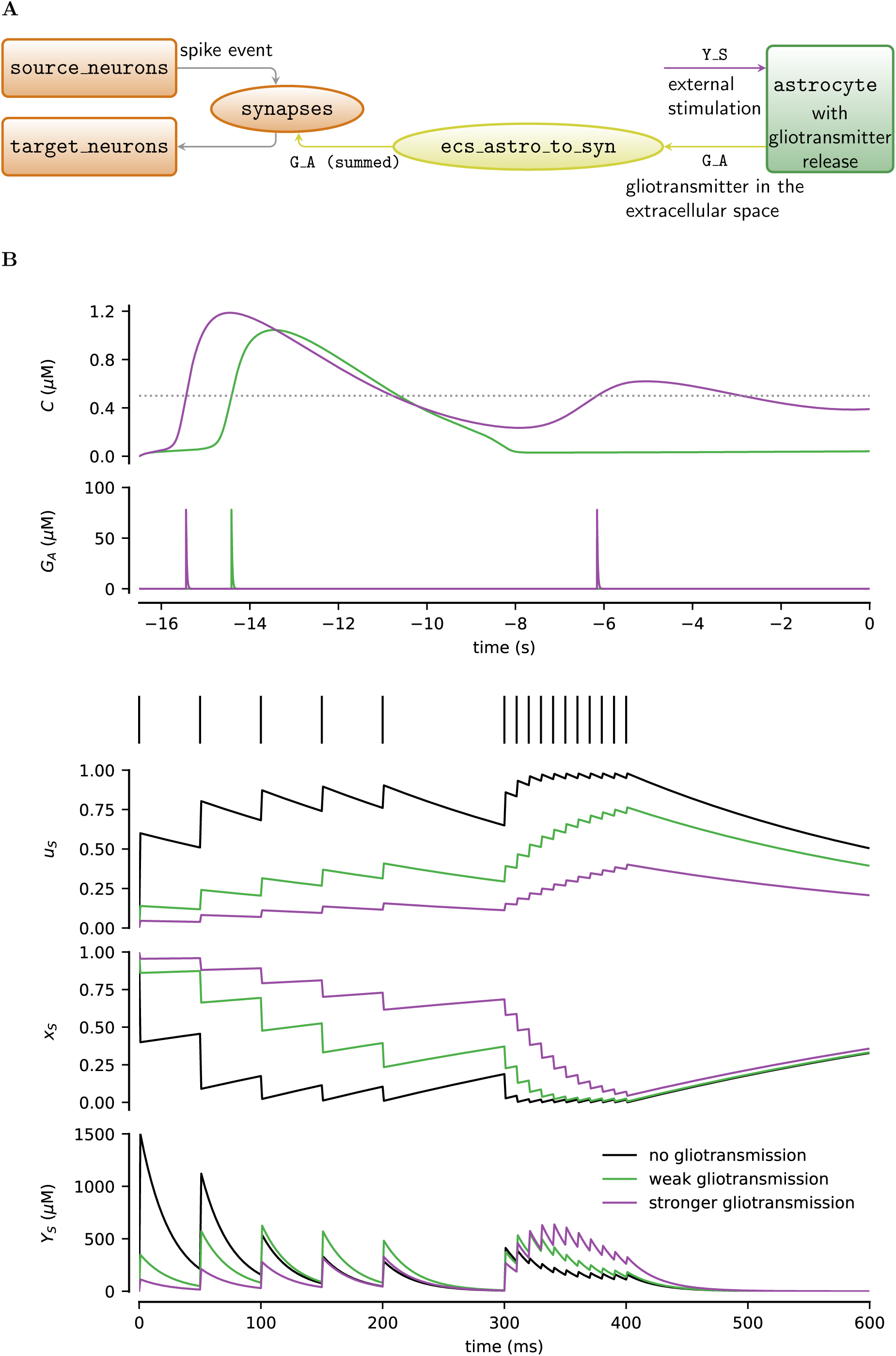
Modeling of modulation of synaptic release by gliotransmission. **A** The model uses a `Synapses` object `ecs_syn_to_astro` to allow synapses’ modulation by perisynaptic gliotransmitters. **B** (*top panels*) Extracellular gliotransmitter concentration (*G_A_*) resulting from intracellular Ca^2+^ dynamics (*C*) in two differently activated astrocytes. (*Bottom panels*) Dynamics of synaptic state variables (*u_S_, x_S_*) and extracellular neurotransmitter concentration (*Y_S_*) in response to a sequence of action potentials (*top*) for a synapse without (*black*) and with (*green* and *purple*) gliotransmitter-mediated modulation. The two astrocytes were initialized at *t* = −17.1s at *x_A_* = 1, *I* = 0.4 μm, and *h* = 0.9 and were respectively stimulated by *I_bias_* = 0.8 μm (weak gliotransmission) and *I_bias_* = 1.25 μm (stronger gliotransmission). Other model parameters as in the Tables in Appendix C.

Figure 3B illustrates how gliotransmitter release from the astrocyte could change synaptic neurotransmitter release in our model. The *top panels* show Ca^2+^ traces (*C*) from two astrocytes that are differently stimulated by exogenous IP_3_ production (*J_ex_* in equation 14) so that their intracellular Ca^2+^ concentration crosses the threshold for gliotransmitter release (*gray dashed line*) at different times. This results in one astrocyte releasing gliotransmitter in the extracellular space (*G_A_*) only once (*green traces*, “weak gliotransmission”), while the other releases gliotransmitter twice (*purple traces*, “stronger gliotransmission”). The modulation of synaptic release ensuing from these two different timings of gliotransmitter release, is illustrated in the *lower panels*, where neurotransmitter release from a synapse stimulated by a sample train of action potentials, is monitored first in the absence of gliotransmitter release from the astrocyte, and then in the presence of weak vs. stronger gliotransmission. Without gliotransmission, the extracellular neurotransmitter concentration (*Y_S_*) progressively decreases with the incoming action potentials, compatibly with the onset of strong short-term synaptic depression (*black traces*). In the presence of gliotransmission instead, while the amount of released neurotransmitter per action potential is generally lower than in the “naive” synapse (since we assumed *α* < 0 in this example), this amount tends to increase at every action potential with respect to preceding ones, and this increase is larger for stronger gliotransmission. This is consistent with the onset of short-term synaptic facilitation, and agrees with the experimental observation and the theoretical argument that gliotransmission could change the synapse’s short-term plasticity (Araque et al., 2014; De Pittà et al., 2016, see also Chapters 8 and 13).

### 2.6 Closed-loop gliotransmission

In the examples discussed so far, we only separately considered one-way interactions between synapses and astrocytes, modeling either modulation of astrocytic activity by synaptic neurotransmitters (Section 2.4) or modulations of synaptic release by astrocytic gliotransmitters (Section 2.5). In practice however, these two pathways may coexist, with gliotransmission feeding back in a closed-loop fashion on the very synapses that stimulate the astrocyte and trigger its gliotransmitter release. This section focuses on such a closed-loop scenario for gliotransmission.

Closed-loop gliotransmission can easily be implemented in *Brian 2* by combining the model of a synaptically-activated astrocyte (Figure 2A) with that of open-loop gliotransmission (Figure 3A), resulting in the model scheme shown in Figure 4A. However, it may be noted that in doing so we end up using two independent `Synapses` objects (*ellipses*) to separately describe the extracellular space for synapse-to-astrocyte signaling `(ecs_syn_to_astro)` and the extracellular space for astrocyte-to-synapse gliotransmission `(ecs_astro_to_syn)`. In reality, both neurotransmitter and gliotransmitter release could occur in the same extracellular space, and thus only a single `Synapses` object might be considered at the benefit of computational efficiency. Nonetheless, the choice of using two independent objects allows us to take into account the more general scenario of astrocytes that are either sensitive or not to the activity of the same synapses they modulate (Martín et al., 2015; Navarrete and Araque, 2010). This is therefore an appropriate choice when dealing with many synapses interacting with astrocytes as in the case of the neuron–glia network discussed at the end of this chapter.

**Figure 4.**
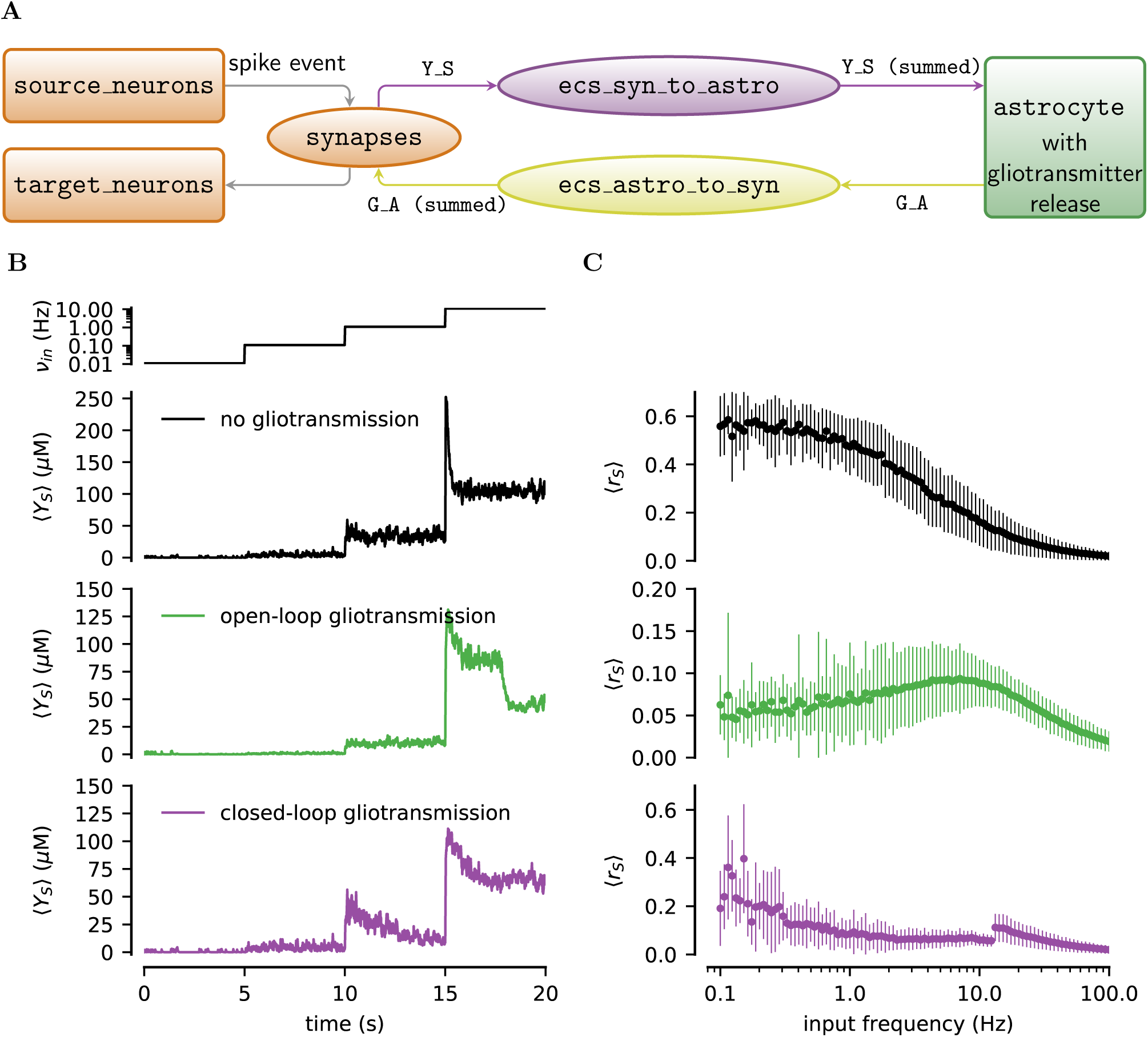
Closed-loop gliotransmission. **A** In the most general design, modeling of closed-loop gliotransmission in *Brian 2* separates between the extracellular space of synapse-to-astrocyte signaling `(ecs_syn_to_astro)` and the extracellular space of astrocyte-to-synapse gliotransmission `(ecs_astro_to_syn)`. **B** Average extracellular concentration of synaptically-released neurotransmitter (*(Y_S_)*) for step increases of the mean rate of Poisson-generated incoming action potentials (*top panel*, *v_in_* = 0.011 Hz, 0.11 Hz, 1.1 Hz, 11 Hz for 5-s time intervals; traces averaged over 500 identical synapses.) **C** Corresponding average release of synaptic neurotransmitter resources as function of the rate of incoming action potentials (data points and error bars: mean *±* standard deviation for 100 trials.) Parameters as in the tables in Appendix C except for *O_β_* = 3.2 μm s^−1^; *I_bias_* = 1 μm (*open-loop gliotransmission*); *I_bias_* = 0 μm (*closed-loop gliotransmission*).

To elucidate some of the possible functional implications of closed-loop gliotransmission we set out to characterize the average synaptic release for `N_synapses=500` identical synapses for different input stimuli and compare it to the open-loop scenario of gliotransmission as well as to the “naive” scenario without gliotransmission. *Brian 2* is optimized to deal with large objects (see Appendix A), so rather than simulating one synapse at a time, for 500 times in the three different scenarios (i.e. 1500 simulations in total), we simulate all synapses in all scenarios in one single run. This is achieved by considering 500 × 3 synapses and an astrocyte group of 500 × 2 elements. The first 500 synapses are modulated by gliotransmitters from the first 500 astrocytes in a closed-loop fashion, the second group of 500 synapses is modulated by open-loop gliotransmission mediated by the other 500 astrocytes; and finally, the remaining synapses do not consider any gliotransmission. In contrast to previous synaptic connection patterns, here we can directly calculate a target index for each connection, instead of evaluating a logical condition for every possible connection pair. *Brian 2* has a built-in syntax for such descriptions, which offers a much more efficient way of establishing connections. In this syntax, we provide an expression to calculate the target index *j* based on the source index *i* and potentially other pre- or postsynaptic properties. To exclude certain potential connections, this expression can be combined with an optional if part stating the condition for a connection to exist. Remember that in our example here, the source index *i* and the target index *j* each refer either to synapses or astrocytes, depending on the direction of the connection (“synapses to astrocytes” or “astrocytes to synapses”). This leads to the following *Brian 2* code:

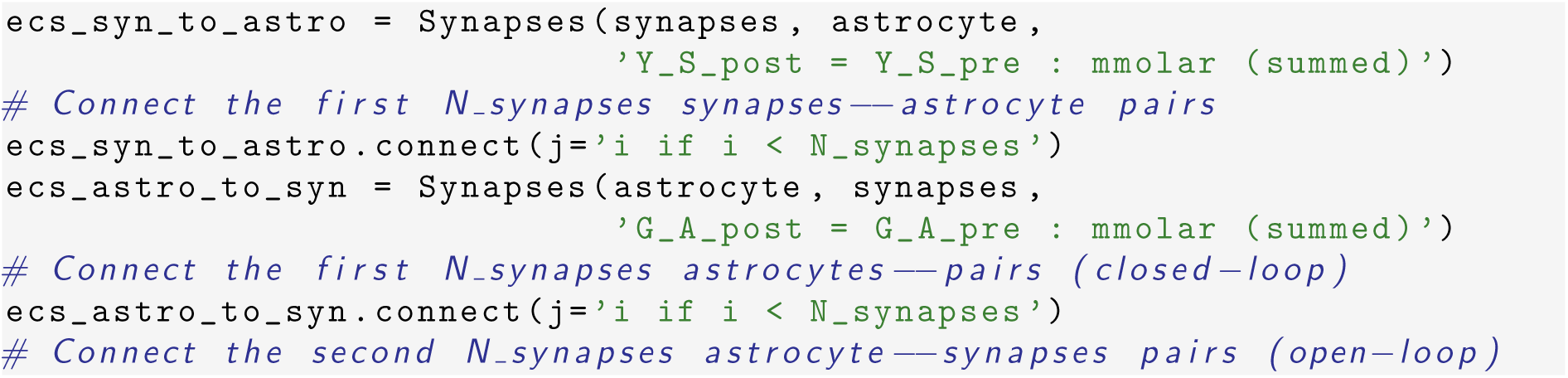

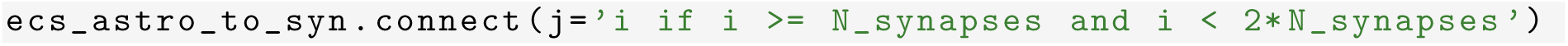

Figure 4B shows a reproduction of Figure 13.2 by our *Brian 2* implementation of closedloop gliotransmission for the time evolution of the average neurotransmitter concentration in the synaptic cleft (*(Y_S_)*) in response to step increases in the rate of incoming action potentials (*v_in_*, *top panel*). Gliotransmission dramatically changes synaptic transmission (*colored* vs. *black traces*), with the effect of closed-loop gliotransmission (*purple trace*) being somewhat intermediate between the scenarios of no gliotransmission (*black trace*) and open-loop gliotransmission (*green trace*).

This is further elucidated in Figure 4C where the mean neurotransmitter concentration in the extracellular space in the three scenarios is shown for different mean rates of randomly incoming action potentials. The low-pass filter characteristics of synapses without gliotransmission (*top panel*) turns into a bell-shaped, band-pass filter characteristics caused by (release-decreasing) open-loop gliotransmission (*middle panel*) (Chapter 12). In the presence of closed-loop gliotransmission however, the average concentration of synaptically-released neurotransmitter is in between those expected in the other two scenarios for low input rate values, and tends to approach the shape of the curve in the open-loop scenario for increasing rates. For high input rates however, the release-decreasing effect of gliotransmission is such that the synapse is ultimately silenced and cannot sustain further gliotransmitter release. Synaptic transmission then becomes independent of gliotransmission again as if it were in the naive scenario without gliotransmission, which accounts for the jump at *v_in_ >* 10 Hz.

### 2.7 Networks of astrocytes

Astrocytes are known to arrange in networks of different shape and connectivity depending on the brain region under consideration (Giaume et al., 2010), and to be capable of propagating Ca^2+^ signals through such networks in the form of intercellular (regenerating) waves. The mechanisms underlying such propagation can be multiple and varied (Scemes and Giaume, 2006). Here, we only focus on the well characterized mechanism of intracellular IP_3_ diffusion through gap junctions channels (GJCs) between neighboring astrocytes (Chapter 7).

From a modeling perspective, IP_3_ diffusion from one astrocyte *j* to a neighboring one *i* can be thought as a flux of IP_3_ (*J_ij_*) which is some nonlinear (rectifying) function of the IP_3_ concentration gradient between cells *i* and *j*, i.e. δ*_ij_I* = *I_i_ − I_j_*, such as, for example (Lallouette et al., 2014, see also Chapter 7)

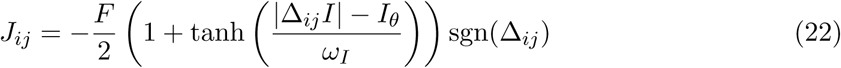

Incidentally, we note that the above formula is reminiscent of the expression of the exogenous IP_3_ flux (*J_ex_*) in equation 14, insofar as the latter may be regarded as a special case of intercellular IP_3_ influx to any astrocyte from a much larger external IP_3_ source (i.e. *I_bias_* in our notation) (Goldberg et al., 2010). Because *J_ij_* is a function of IP_3_ concentrations in connected astrocytes (i.e. *I_i_, I_j_*) by δ*_ij_I*, it is astrocyte-dependent and not constant. Therefore, once we add *J_ij_* to our astrocyte equations in *Brian 2* (denoted in the code below by `J_coupling`), we must define it as an astrocytic variable (that is without the (constant) flag), i.e.

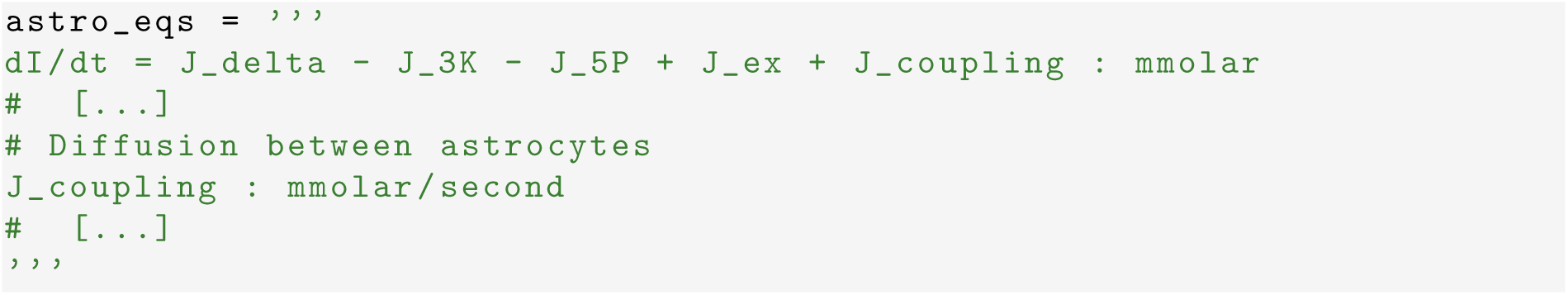

Connections between astrocytes by GJCs may conveniently be implemented by a `Synapses` object in *Brian 2*, once we regard *J_ij_* as the IP_3_ flow from “presynaptic” astrocyte *j* to “postsynaptic” astrocyte *i* (Figure 5B). The effective total `J_coupling` to cell *i* by intercellular IP_3_ diffusion is the sum of all IP_3_ fluxes incoming to cell *i* from the *A*^*i*^ astrocytes connected to this latter by GJCs, i.e. 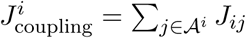. In *Brian 2* code, this reads

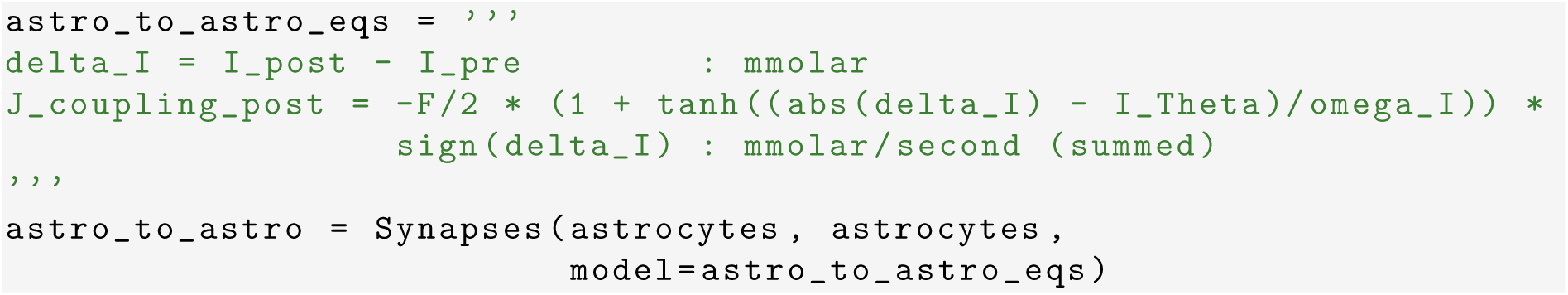

**Figure 5.**
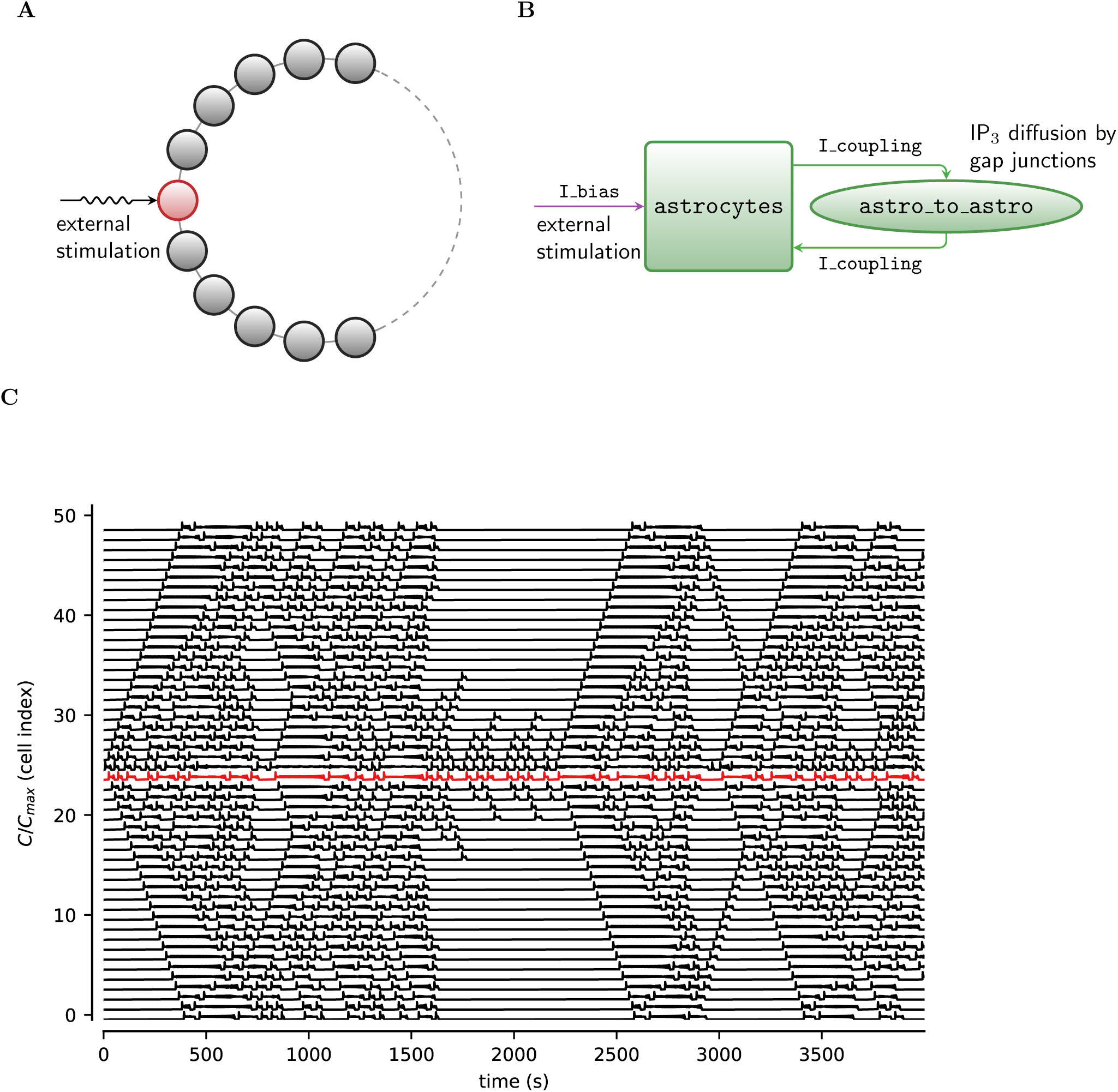
Astrocytes connected in a network. **A** Sample astrocyte network in a ring configuration with only one cell (in *red*) being exogenously stimulated. Connections between cells are bidirectional, and represent GJC-mediated coupling between neighboring astrocytes. **B** General *Brian 2* modeling principle of astrocytic networks: GJC-mediated connections can be modeled by a `Synapses` object (*ellipse*). **C** Intercellular Ca^2+^ wave generation and propagation in a ring of 50 identical astrocytes mediated by stimulation of cell 25 (*red trace*). Parameters as in Table C.2 with *F_ex_* = 0.09 μm s^−1^; and *I_bias_* = 1 μm.

The above code bears the caveat of defining GJCs as unidirectional when they may not be so. This caveat can be easily overcome, specifying both a connection from astrocyte *i* to astrocyte *j* and a connection from *j* to *i*, whenever we want to model bidirectional IP_3_ diffusion between neighboring astrocytes. For example, to connect astrocytes in a ring, where every astrocyte is connected to its neighbors (Figure 5A), we can make use of the connect method of the `astro_to_astro` object, and specify the following condition in terms of *Brian 2* predefined pre- and post-synatic indexes, i and j respectively, and the total number of elements in the presynaptic group N_pre^3^:

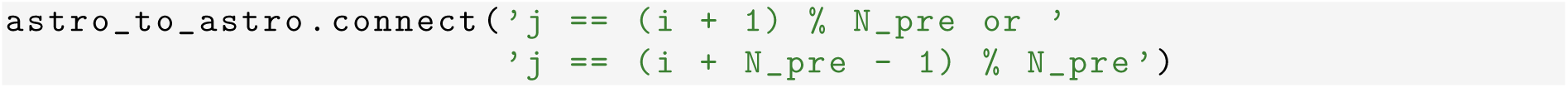

where the % operator implements the modulo (remainder) operation.

Figure 5C shows a snapshot of Ca^2+^ dynamics of 50 astrocytes connected in a ring, where only the 25th cell is exogenously stimulated (*red trace*). The fact that all cells, for some *t* > 0, display Ca^2+^ fluctuations, is a direct consequence of inclusion of intercellular IP_3_ diffusion in our model. Such diffusion allows excess IP_3_ from the stimulated cell to be redistributed by GJCs in the ring to other cells where it ultimately triggers CICR. It may also be appreciated how, in this example, bidirectional GJC communication allows for emergence of intercellular Ca^2+^ waves that propagate both from and to the stimulated cell, as evidenced by wave fronts respectively oriented like ‘*\*’ or like ‘/’.

### 2.8 Coupled neuron and astrocyte networks

The examples discussed so far provide together all the ingredients to model complex networks of interacting neurons and astrocytes (Figure 6A). However, to realistically implement such networks we also need to specify the connections among neurons, synapses and astrocytes in the physical (Euclidean) space. In the following we show how to include space in such networks, limiting our focus here to planar networks for simplicity, although the outlined procedure can easily be extended to higher dimensions.

**Figure 6.**
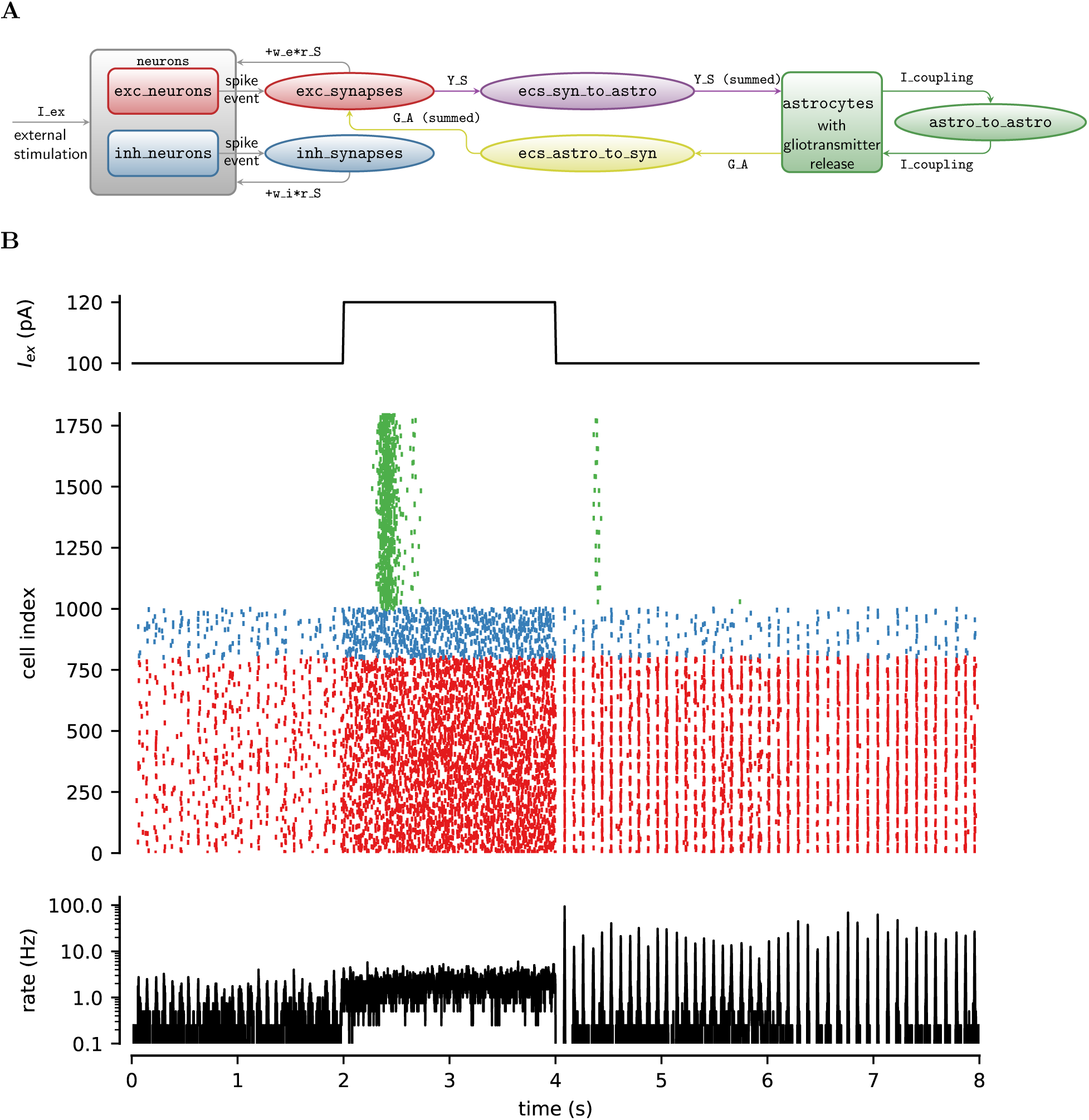
Recurrent neuron-glial network. **A** Neuron-glial network model design in *Brian 2*. **B** Simulations of neuron-glia network for a rectangular-pulse increase of external current (*I_ex_*, *top panel*). The raster plot (*middle panel*) shows the firing activity of 25% out of all excitatory (*red*) and inhibitory neurons (*blue*) of the network, and gliotransmitter release (*green*) from an equal fraction of astrocytes. The network-averaged firing rate is shown at the *bottom*. Neural activity dramatically changes from asynchronous low-firing activity to synchronous high-firing activity following gliotransmitter release from astrocytes during the period of high stimulation (2 ≤ *t* < 4 s). External current: *I_ex_* = 100 pA for *t* < 2s or *t* ≥ 4 s; *I_ex_* = 120 pA for 2 ≤ *t* < 4 s. Neural and synaptic parameters as those in Figure 1 (see also Table C.1). Astrocyte parameters as in Table C.2 except for *O_β_* = 0.5 μm s^−1^; *O_δ_* = 1.2 μm s^−1^; and *I_bias_* = 0.

We start by adding two cell-specific parameters, *x* and *y*, to each neuron which store the cell’s 2D spatial coordinates and initialize them so that neurons are arranged on a grid of `N_rows rows` and `N_cols columns`:

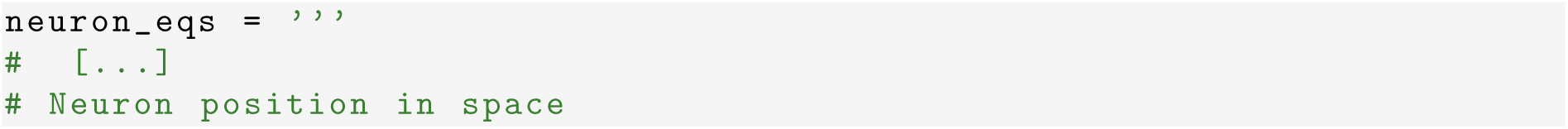

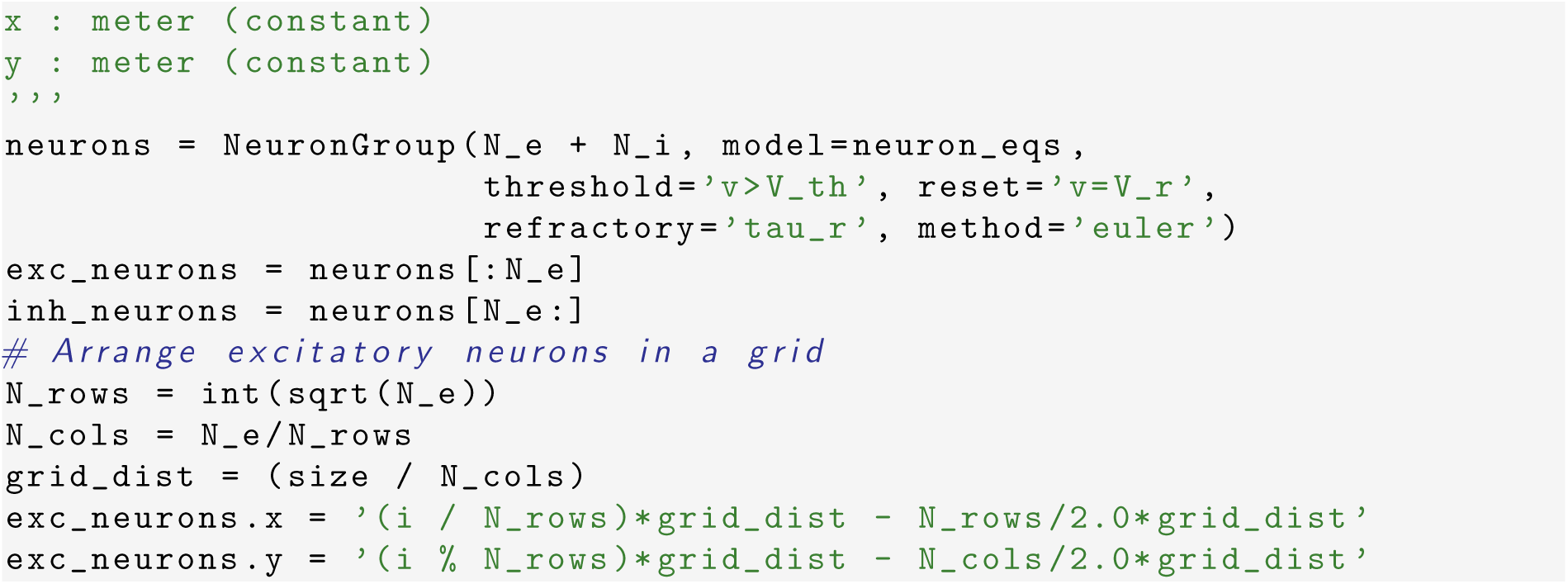

Furthermore, we also add a synapse-specific constant `astrocyte_index` to the synapse’s equations, whose value will correspond to the index of the astrocyte that ensheathes a synapse:

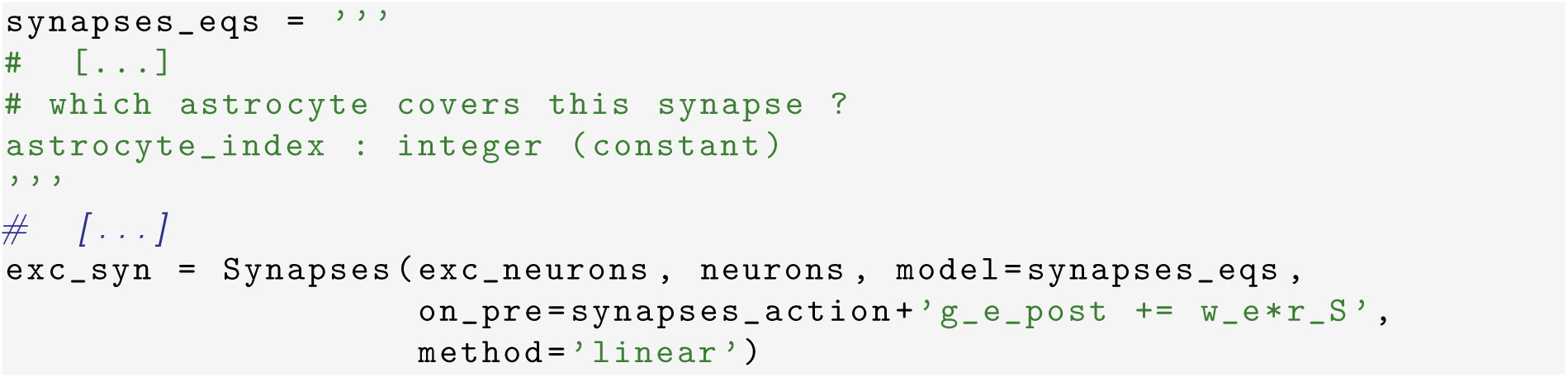

We finally need to define the effective connections between the different cells of the network. Overall there are four different types of connections: (i) connections between neurons which defines the actual synapses; (ii) connections from synapses to astrocytes, as pathways to trigger astrocyte activation; (iii) connections from astrocytes to synapses as routes for gliotransmission and thereby modulation of synaptic release; and ultimately, (iv) connections between astrocytes by GJCs. Here, for simplicity, we assume random connectivity between all neurons, independently of their spatial coordinates (as in Figure 1C). Furthermore, we make the assumption that only excitatory synapses can activate astrocytes and be modulated by them, restricting in this way our focus on the experimentally well-characterized pathway of closed-loop glutamatergic gliotransmission (Panatier et al., 2011; Perea and Araque, 2007). In particular, we specify which astrocyte is responsible for which excitatory synapse on the basis of the spatial position of postsynaptic neurons with respect to *N_a_* astrocytes `(N_a)` which, like neurons, are arranged on a regularly-spaced grid of *N_rows_* rows `(N_rows_a)` and *N_cols_* columns `(N_cols_a)`, i.e.

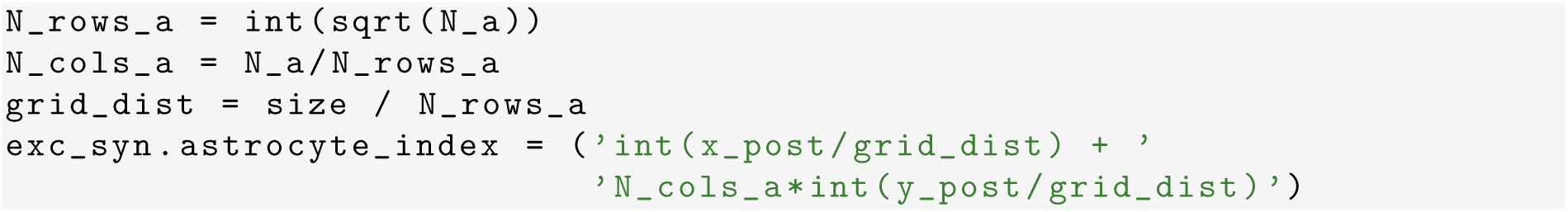

We then define the network of astrocytes:

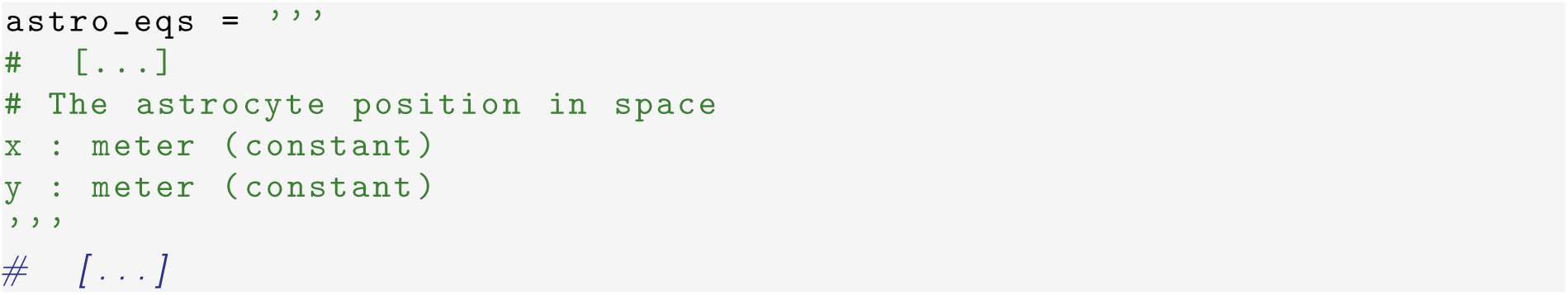

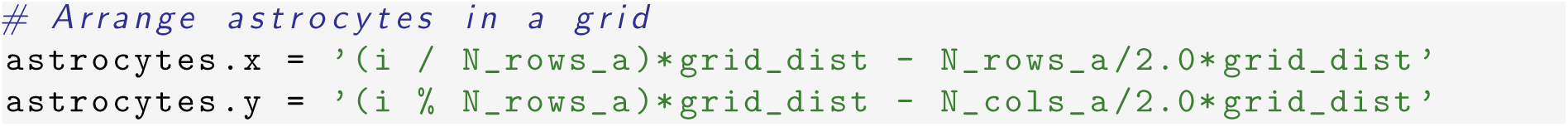

Next, we connect the astrocytes with those synapses that they are supposed to ensheathe according to astrocyte_index, i.e.

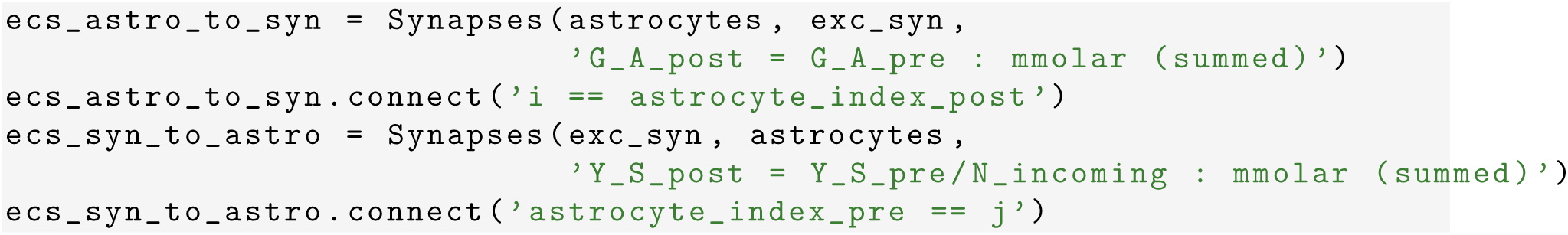

Finally, we specify the connectivity of the astrocyte network. In this example, we introduce recurrent connections between astrocytes by GJCs, connecting each astrocyte to all other astrocytes found at the boundary of its anatomical domain, in line with the experimental observation that neighboring astrocytes are more likely to be connected than astrocytes that are far apart (Giaume et al., 2010; Pannasch and Rouach, 2013). Given that the diameter of astrocyte is between 50–130 μm (Chao et al., 2002), we consider an intermediate value of 75 μm, whereby:

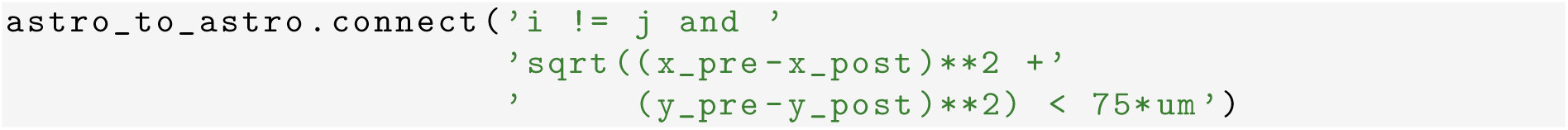

We present a simulation of our neuron-glia network in Figure 6B, where we show the raster plot of the firing activity of 25% of the excitatory (*red*) and inhibitory neurons (*blue*) of the network along with gliotransmitter release events from an equal fraction of astrocytes (*green*), in response to a transient increase of external stimulation (rectangular pulse in the *top panel*). Up to the onset of stimulation (i.e. *t* < 2 s) there is no gliotransmitter release from astrocytes, therefore the network behaves as it would be expected for a neuronal network without the astrocyte component. It may be noted in fact how the raster plot of our network, and the ensuing dynamics of the total firing rate (*bottom panel*), show low-frequency population activity, similar to those reported in Figure 1C for our neuronal-only network model introduced in Sections 2.2 and 2.3. For 2 ≤ *t* < 4 s, the increase of external stimulation correlates with an increase in the firing rate of the whole network, as reflected by a denser raster plot during this period. In particular for *t>* 3.5 s, the larger neuronal firing triggers gliotransmitter release from astrocytes and thus astrocytic modulation of excitatory synaptic transmission. Because this modulation is slow-decaying (Chapter 8), it outlasts the transient increase of external stimulation and changes neural firing once the external stimulation returns to its original value (at *t* = 4 s). We can indeed clearly see how, for *t >* 4 s, excitatory neurons are more synchronized in firing than for *t <* 2 s, as a consequence of gliotransmission from astrocytes. This is just one example of the many possible ways astrocytes could actively shape neural activity, which has also been suggested to participate in the genesis of cortical UP and DOWN states (Fellin et al., 2012).

## Conclusions

Computational approaches to model glial physiology are hampered by the lack of definitive experimental evidence and a missing comprehensive modeling framework that could tackle the many different scales of glial signaling. “Standard” glia models have yet to be identified, and neural simulator packages therefore do not ship such models as part of their pre-built model libraries. While in theory these libraries could be extended by individual researchers to add their preferred glia model, in practice this path is only open to experienced programmers.

In this chapter, we have shown how *Brian 2*’s simple syntax and versatility can offer a solution to these problems, providing an ideal tool to model glial physiology, and specifically the influence of astrocytes on neural activity. *Brian 2*’s syntax allows the researcher to flexibly describe models by using conventional mathematical notation instead of low-level programming code (Goodman and Brette, 2008; Goodman et al., 2009; Stimberg et al., 2014). Moreover, *Brian 2*’s core data structure NeuronGroup, which describes a neuron by a set of ODEs, parameters, and actions that are triggered by conditions, provides a versatile framework that can be borrowed to also describe non-neuronal cell types such as astrocytes. Similarly, the `Synapses` data structure that, in purely neural simulations, represents chemical and electrical synapses that connect neurons, can also be used to model the interactions between astrocytes and synapses, as well as GJCs between astrocytes. Importantly, this flexibility does not come at the cost of computational efficiency: without any user interaction, *Brian 2* employs a code generation approach that generates highly efficient code based on the user-provided high-level description (Goodman, 2010). We hope that these arguments motivate newcomers as well as experienced researchers to experiment with *Brian 2* in the future and use it to model glial physiology in their research, thereby contributing to the growth of this exciting emerging field of computational research.

## Appendix A Technical remarks on *Brian 2*

*Brian 2* scripts are executed by default in the so-called “runtime mode”. This mode runs the simulation loop over the time steps in Python and executes chunks of target language code that have been generated from the model description provided by the user. The choice of target language depends on the user’s system; *Brian 2* will prefer to use the C++ programming language but, if the user does not have a working C++ compiler, will fall back to a pure Python-based simulation. A Python-based simulation will usually be significantly slower but can give comparable performance for big networks due to the use of vectorized computation (Brette and Goodman, 2011). The advantage of the runtime mode is that the user has full control to combine the automatically generated simulation code with arbitrary hand-written Python code. This code could dynamically change aspects of the model during the run, or interact with it in other ways. For example it could read out the model’s state and hand it over to some code for visualization or terminate the simulation based on some criterion. This mode however involves a significant overhead per simulated time step, since the program flow constantly switches between Python and the individually-generated code chunks. For small-to-medium size networks for which computations during a single time step do not take long, this overhead can critically dominate the total runtime and lead to long simulation times.

To avoid this problem and allow more efficient simulations, *Brian 2* also offers an alternative mode called “standalone mode”. In this mode, the complete simulation code, including the main simulation loop, are written as a set of C++ files to disk which can then be compiled and executed as a single program. The resulting files are independent of the Python platform, so that the simulation could also be run on systems where Python may not be available (for example, in robotics). Moreover, if the user code complies to some specific conventions and does not run custom Python code during a simulation, then switching from runtime to standalone mode only requires the addition of a `set_device(’cpp_standalone’)` line to the simulation script; *Brian 2* then takes care of the whole process transparently. For further details, the reader is invited to see comments in individual examples files (Appendix B) and/or refer to the online *Brian 2* documentation.

## Appendix B Example files

The code for all the simulations presented in this chapter has been organized in multiple standalone example files as detailed in the following. Unless stated otherwise, all simulations start from zero initial conditions, except for *h*(0) = 0.9 and *x*_*S*_ (0) = *x_A_*(0) = 1.

~~~
example_1_COBA.py
~~~

This file implements the simulation of the neuron-only network model of Figure 1. The simulation runs for 1 s with an integration time step of 0.1 ms. Out of all neurons, we distinguish between excitatory `(exc_neurons)` and inhibitory ones `(inh_neurons)`, which give rise to excitatory synapses (exc_syn) and inhibitory synapses (inh_syn), connecting from the respective population to the full population. Because the dynamics of synaptic variables are updated only at incoming action potentials (i.e. (event-driven)), we can monitor the value of these variables only at the arrival time of action potentials but not in between. However, we can reconstruct the whole synaptic dynamics by recording synaptic variables immediately after each action potential (i.e. at 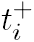 with *i >* 0), which is achieved by specifying the keyword argument when=’after_synapses’ in the synaptic StateMonitor. For *t* > *t_i_*, the solutions of the synapse’s equations 4 and 5 then read (Tsodyks, 2005):

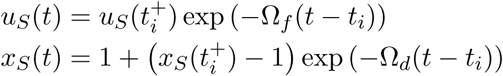

whereas synaptic release by the *i*th action potential at time *t_i_* is given by 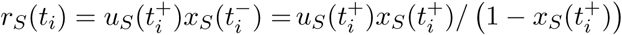

~~~
example_2_gchi_astrocyte.py
~~~

This code implements the synaptically-stimulated astrocyte model and related simulations of Figure 2. The astrocyte’s temporal dynamics in response to synaptic activity was simulated for 30 s using the derivative-free Milstein integration method with a time step of 1 ms. In the deterministic limit of *ξ*(*t*) *→* 0 in equation 13, the Milstein method reduces to the classical (forward) Euler method which is suitable, at sufficiently small time steps, to numerically solve dynamics of the deterministic astrocyte model, too. `Synapses` are stimulated by a train of periodic action potentials at rate *f*_0_ = 0.5 Hz (f_0, rate of generation of action potentials by presynaptic neurons) generated by

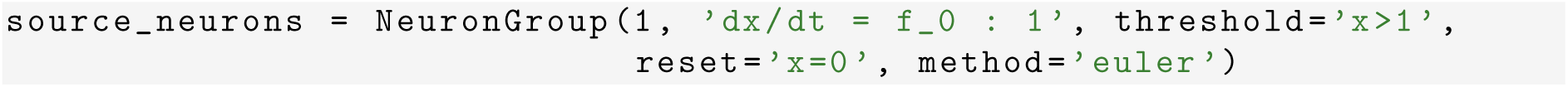

~~~
example_3_io_synapse.py
~~~

This file implements the open-loop model of gliotransmission and the simulations shown in Figure 3. The code considers three synaptic connection between one presynaptic source_neurons and one postynaptic target_neurons, built by passing n=3 as an argument to the synapses.connect method. We further consider two astrocytes stimulated by different `I_bias` values, and connect them to synapses 2 and 3 respectively, leaving synapse 1 as it is (i.e. without gliotransmission). This is done by:

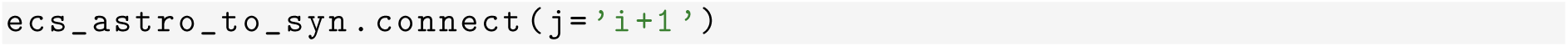

~~~
example_4_synrel.py
~~~

This code runs the closed-loop model of gliotransmission for simulations in Figure 4B. The code considers N_synapses neurons `(source_neurons)`, each firing action potentials drawn from an independent, inhomogeneous Poisson process with a stepped rate specified in a `TimedArray`, i.e.

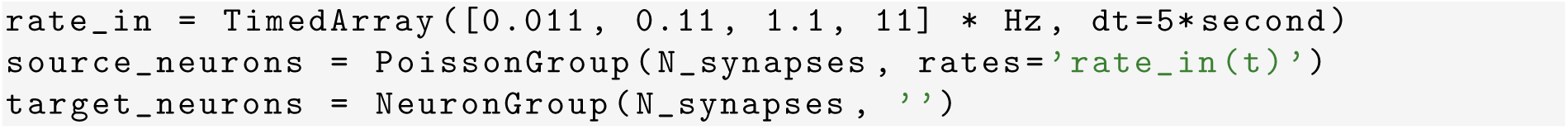

The `target_neurons` are used to build `N_synapses` multi-synaptic connections from the `source_neurons`, with each connection constituted of three synapses. Out of these three synapses, the first one is connected with its own astrocyte and is, in turn, modulated by gliotransmitters released from this latter (closed-loop scenario); the second one is modulated by gliotransmitters released from another astrocyte (open-loop scenario); the third one is left as it is (scenario without gliotransmission). Since this is repeated for all `N_synapses`, and overall we have `N_astro=2` different scenarios of gliotransmission (open-loop vs. closed-loop), we consider `N_astro*N_synapses` astrocytes in total, and connect them accordingly with `N_synapses*(N_astro+1)` synapses as elucidated in Section 2.6.

~~~
example_4_rsmean.py
~~~

The file provides the code to build the synaptic transfer characteristics in Figure 4C in terms of average synaptically-released neurotransmitter resources for different input rates of (presynaptically) incoming action potentials.

~~~
example_5_astro_ring.py
~~~

This code implements the astrocyte ring model in Figure 5. The simulation runs for 4000 s with a time step of 50 ms. Calcium concentrations shown in Figure 5C were normalized by their maximum.

~~~
example_6_COBA_with_astro.py
~~~

This file runs the simulation of the recurrent neuron-glial network in Figure 6. To stimulate the network by a time-varying external current we multiply I_ex in `neuron_eqs` on page 4 by stimulus = TimedArray([1.0, 1.2, 1.0, 1.0], dt=2*second). Neurons are placed on a square lattice of size 3.75 × 3.75 mm at 50 μm distance from each other. For *t* = 0 we set *C* = *I* = 0.01 μm.

## Appendix C Model parameters used in the simulations

The following tables report constants that correspond to the model parameters used in the simulations presented in this chapter. Simulation-specific parameters are marked by ‘*†*’ and are reported in respective figure captions instead.

## Footnotes

1 Note that molar is *not* a SI base unit, because it is defined as m = mol L^−1^, i.e. referring to L instead of the SI base unit m^3^. Since 1 mol L^−1^ = 1000 mol m^−3^, the base unit to use is mm (mmolar).

2 *Brian 2* also offers an alternative system to specify constants via a namespace argument that receives a Python dictionary mapping constant names to their values. Refer to *Brian 2*’s online documentation for details at http://brian2.readthedocs.io.

3 Note that the expression has been split into two strings for better readability. Python automatically merges adjacent strings.

